# Visualization of silver nanoparticle intracellular trafficking revealed nuclear translocation of silver ions leading to nuclear receptor impairment

**DOI:** 10.1101/825919

**Authors:** Vanessa Tardillo Suárez, Elizaveta Karepina, Mireille Chevallet, Benoit Gallet, Cécile Cottet-Rousselle, Peggy Charbonnier, Christine Moriscot, Isabelle Michaud-Soret, Wojciech Bal, Alexandra Fuchs, Rémi Tucoulou, Pierre-Henri Jouneau, Giulia Veronesi, Aurélien Deniaud

## Abstract

The impact on human health of the increasing use of silver nanoparticles (AgNPs) in medical devices remains understudied, even though AgNP-containing dressings are known to release silver in the bloodstream leading to accumulation and slow clearance in the liver. Cellular studies have shown the intracellular dissolution of AgNPs within endo-lysosomes followed by Ag(I) binding to biomolecular thiolate-containing molecules. However, the precise subcellular distribution of Ag(I) and the nature of the disrupted physiological pathways remained unknown. Novel imaging approaches enabled us to visualize the trafficking of AgNP-containing lysosomes towards a perinuclear location and a direct nuclear transfer of Ag(I) species with accumulation in the nucleoli. These Ag(I) species impaired nuclear receptor activity, disrupting critical mechanisms of liver physiology in very low dose exposure scenarios, thus justifying further research into defining a framework for the safe use of AgNPs.

## 1. Introduction

Silver nanoparticles (AgNPs) are widely used in everyday products for their biocidal activity and increasing production has led to environmental and human safety concerns over the past few years. Their biocidal properties stem from the durable release of Ag(I) ions from the NP’s Ag(0) core due to surface oxidation, ions that are also toxic for mammalian cells.

In humans, the main and most extensively described exposure route for AgNPs is dietary.^1, 2^ However, the impact of nanosilver-containing medical devices that can directly release Ag into lymphatic and blood vessels has been sorely disregarded. The comparison of Ag distribution and excretion after a single oral or intravenous exposure to AgNPs in mice showed a five to ten times higher blood concentration after 24 hours in the latter case.^3^ In both cases, Ag preferentially accumulated in the liver but Ag clearance was much slower when AgNPs were injected. This is most probably due to the presence of Ag in the NP form mainly as Ag(0), whereas during oral exposure AgNPs are readily transformed in the gastrointestinal tract, and Ag enters the bloodstream in the form of various Ag(I) species (^4^ and for a recent review see^1^). Therefore, it is of paramount importance to better understand AgNP fate in the liver to predict the risk for patients during hospital care. In spite of its relevance, this exposure scenario has been as yet disregarded in nanotoxicology studies. In this context, we decided to address the impact of an exposure over several days to low and non-toxic concentrations of pristine AgNPs on critical hepatocyte functions.

Over the past few years, several studies have described the fate of AgNPs in various mammalian cell models.^5–9^ A consensus has emerged showing AgNP dissolution in endosomes and lysosomes and the subsequent distribution of Ag(I) species throughout the cell mainly in the form of Ag-thiolate complexes.^6, 8–10^ This type of complexation comes as no surprise as Ag has a picomolar or higher affinity for thiolates depending on the coordination chemistry offered by the biomolecular ligands.^11^ However, recent imaging studies did not provide the subcellular distribution of Ag(I) species at the organelle level.^9, 12^ The identification of cell compartments that can store Ag and/or that can be critically affected by this non-physiological and deleterious metal ion remains a critical question. Of particular importance is the long-standing controversy about AgNP entry into the nucleus^13, 14^ (for a recent review see^1^).

In the current study, we developed a methodology which allowed us to map the distribution of any form of Ag, even ionic, at the level of the organelle and combined it with other imaging approaches. These experiments were carried out under long-term low dose exposure in order to be as realistic as possible. We thus highlighted that Ag(I) species but not AgNPs are translocated to the nucleus and we further analyzed their impact on nuclear functions. Hepatocyte nuclear receptors, such as the Liver X receptor (LXR) and the Farnesoid X receptor (FXR), also known as the bile acid receptor (BAR), were identified as molecular targets that were inhibited by these translocated metal ions even under a very low dose exposure scenario mimicking chronic exposure. AgNPs could thus trigger an endocrine disruptor-like effect.

## 2. Experimental

### 2.1. AgNP characterization

Highly homogenous 20 nm citrate-coated AgNPs were purchased from NanoComposix (20 nm Citrate BioPure^™^ Silver) and 90 nm PVP-coated AgNPs were purchased from Sigma-Aldrich.^9^ Dynamic light scattering measurements were carried out in cell culture medium supplemented with fetal calf serum at 25 °C in a UVette (Eppendorf) on a Nanostar instrument (Wyatt). Zeta potential were determined using a Malvern Nano ZS in 10 mM HEPES, 2 mM sodium citrate pH 7. In cell culture media, citrate- and PVP-coated AgNPs formed agglomerates of 160 nm and homogenous 90 nm particles, respectively (Table S1).

### 2.2. Cell culture

HepG2 cells were grown in MEM media and exposed to cit-AgNPs or PVP-AgNPs at the indicated concentrations of total Ag and for the indicated times. Media and AgNPs were renewed every 24 hours. When required, chenodeoxycholic acid (CDCA, Sigma-Aldrich) at 80 μM or the equivalent volume of DMSO was added for the last 17 hours of exposure or 22R-hydroxycholesterol (22R-HC, Sigma-Aldrich) at 10 μM or the equivalent volume of absolute ethanol was added for the last 20 hours of exposure.

### 2.3. Spheroid preparation

Micropatterns were produced on glass slides by deep-UV photolithography by which a non-toxic cell repellent PEG-PLL coating was degraded locally by deep-UV light passing through a photomask (as described in^15^). Micropatterns were then coated with fibronectin (20 μg.mL^-1^ in PBS) to form adhesive disks of 50 μm in diameter, surrounded by a cytophobic surface.

HepG2/C3a cells were seeded on micropatterned surfaces at a density of 1.25 10^5^ cells per cm^2^ and were allowed to attach. After 2 hours, unattached and floating cells were removed by extensively washing the surface and gentle flushing with a pipette. Cells on micropatterns were maintained in culture for up to one week, to allow spheroids to grow.

### 2.4. Subcellular fractionation and elemental quantification

Subcellular fractionation was performed using a protocol developed to purify nucleoli^16^ that we adapted to avoid the contamination of the nuclear fraction by endo-lysosomal vesicles rich in AgNPs. Indeed, at low centrifugation speed, the high density of trapped AgNPs pelleted the “light vesicles” containing AgNPs. Therefore, we purified nuclei through a discontinuous sucrose/percoll gradient centrifugation in which nuclei are recovered at the interface between two layers and not at the bottom of the tubes where AgNP-containing vesicles can be found.

After AgNP exposure for the indicated times, HepG2 cells were recovered by trypsinization. The cell pellet was resuspended in 7.5 mL of 10 mM Tris-HCl pH 7.4, 10 mM NaCl and 1 mM MgSO4 and incubated for 20 minutes on ice with regular mixing. NP40 was added at a final concentration of 0.3% and cell lysis was carried out with a Dounce homogeneizer. The nuclei-containing pellet was collected by centrifugation at 1,200 g for 5 minutes. The pellet was resuspended in 2.5 mL of 10 mM Hepes pH 7.5, 10 mM MgSO_4_ and 0.25 M sucrose and loaded on top of a gradient of 10 mM Hepes pH 7.5, 0.25 M sucrose and 0.05 mM MgSO_4_ with a 2.5 mL layer of 20% percoll, a 2.5 mL layer of 50% percoll, a 2.5 mL layer of 60% percoll and a 1 mL layer of 70% percoll. The gradient was centrifuged for 1 hour at 100,000 g and 4 °C. The main fraction, the nuclei, was recovered at the interface between the 20 and 50% percoll layers and diluted in 10 mL of 10 mM Hepes pH 7.5, 0.25 M sucrose and 10 mM MgSO_4_. This fraction was centrifuged 10 minutes at 1,200 g through a layer of 10 mL of 10 mM Hepes pH 7.5, 0.88 M sucrose and 0.05 mM MgSO_4_. The pellet contained pure nuclei that were resuspended in 10 mL of 10 mM Hepes pH 7.5, 0.34 M sucrose and 0.05 mM MgSO_4_. A fraction was kept for further analysis and the main part was sonicated 4 times 30 seconds at 10% power with a microtip (Sonicator BioBlock Scientific Vibracell 72412). This solution was centrifuged 20 minutes at 2,000 g through a layer of 10 mL of 10 mM Hepes pH 7.5, 0.88 M sucrose and 0.05 mM MgSO_4_. The nucleolar pellet was resuspended in 100 μL of 10 mM Hepes pH 7.5, 0.34 M sucrose and 0.05 mM MgSO_4_. The protein content of the nuclear and nucleolar fractions was quantified using the microBCA method (Pierce) following the manufacturer’s instructions.

The amount of Ag present in the cells and in nuclei and nucleoli fractions was quantified by inductively coupled plasma atomic emission spectroscopy (ICP-AES) using a Shimadzu ICPE-9800 after an overnight mineralization at 50 °C in 65% HNO_3_.

### 2.5. RNA extraction and quantification

At the indicated timepoints, cells were harvested and mRNA were isolated using the Nucleospin RNA kit (Macherey-Nagel) for 2D cultures or the NucleoSpin RNA Plus XS kit (Macherey-Nagel) for the spheroid samples. The RNA concentration was determined using a NanoDrop spectrophotometer (ND-1000). RNA were reverse transcribed with the Affinity script qPCR cDNA synthesis kit (Agilent), according to the manufacturer’s instructions using random primers. Quantitative PCR was performed as described in,^9^ qPCR reactions were run in triplicate, and quantification was performed using comparative regression (Cq determination mode) using Cfx (Bio-Rad Cfx manager) with GAPDH and HPRT amplification signals as housekeeping genes to correct for total RNA content, while the untreated sample was used as the “calibrator”. At least three biological replicates were performed for the presented data. The primers used are described in Table S2, ESI†.

We performed the analysis of FXR and LXR activities based on the induction of their target genes, *bsep* (coding for Bile Salt Export Pump, BSEP) and *abcg5* (coding for the ATP-binding cassette subfamily G member 5), respectively. In this case, the control (ctl) sample (unexposed to AgNPs) was set to 100% activity of FXR or LXR and it corresponds to the ratio:

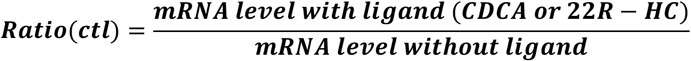

For all conditions of AgNP exposure, we similarly calculated the ratio:

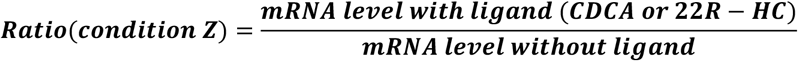

In order to determine the percentage of activity in condition Z, we applied the following equation:

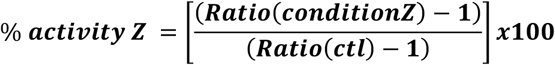

Subtracting 1 takes into account that the value of a control in mRNA normalized relative expression is 1.

### 2.6. Sample preparation for EM and Nano-XRF

At the desired timepoint, monolayers of HepG2 cells were fixed overnight at room temperature in a 1:1 ratio mixture of 4% paraformaldehyde, 0.4% glutaraldehyde in 0.2 M PHEM pH 7.2 and culture medium, washed in 0.1 M PHEM pH 7.2, and fixed for 30 minutes in 2% paraformaldehyde, 0.2% glutaraldehyde in 0.1 M PHEM pH 7.2, washed in 0.1 M PHEM pH 7.2 and post-fixed in 1% OsO_4_ in 0.1 M PHEM buffer for 1 h at room temperature. Cells were then dehydrated in graded ethanol series, and flat-embedded using an Epoxy Embedding Medium kit (Sigma-Aldrich). Sections (200 or 400 nm) were cut on a Leica UC7 ultra-microtome using a DiATOME 35° diamond knife and collected on 50 nm-thick Si_3_N_4_ grids (Oxford Instruments). The same preparation protocol was used for spheroids, where some sections were collected on formvar carbon coated 100 mesh copper grids for scanning transmission electron microscopy coupled with energy-dispersive X-ray spectroscopy (STEM-EDX) and the block was then used for focused ion beam scanning electron microscopy (FIB-SEM).

### 2.7. Nano-XRF data acquisition and analysis

X-ray fluorescence (XRF) experiments were carried out at the state-of-the-art hard X-ray nanoprobe beamline ID16B at the ESRF.^17^ The incoming photon energy was set at 29.6 keV and the beam was focused using Kirkpatrick-Baez mirrors down to 50×50 nm^2^. The fluorescence emission from the sample was recorded using two 3-element Silicon Drift Detectors (SDD) arrays positioned at 13° from the sample. The photon flux on the sample was ~10^11^ photons/s. High resolution maps were recorded at room temperature, raster scanning the sample in the X-ray focal plane with 100×100 nm^2^ step size and 500 ms dwell time/pixel.

Hyperspectral images were analyzed using the PyMCA software package (http://pymca.sourceforge.net/).^18^ The detector response was calibrated over a thin film multilayer sample from AXO (RF8-200-S2453). XRF data were energy calibrated, normalized by the incoming photon flux, and batch-fitted in order to extract spatially resolved elemental concentrations, assuming a biological matrix of light elements and density 1 g.cm^-3^ according to NIST standards (https://physics.nist.gov/cgi-bin/Star/compos.pl?matno=261). At least three different cells per exposure condition were analyzed, and their averages and standard deviations are reported in Fig. 3A.

### 2.8. X-ray absorption spectroscopy (XAS) data acquisition and analysis

Ag K-edge X-ray absorption spectra were acquired on the bending-magnet beamline BM23 of the ESRF.^19^ HepG2 cells were exposed to AgNPs for the indicated times, pelleted and resuspended in PBS containing 20% glycerol (20 μL). Drops of cell suspensions or reference solutions (50 μL) were deposited in the XAS sample holder equipped with Kapton windows and immediately frozen in liquid N_2_, then transferred into the He cryostat and measured at 10 K. The incoming photon energy was scanned around the Ag K edge (25.514 keV) with a fixed exit double crystal Si(311) monochromator (Kohzu, Japan) cooled with liquid N_2_. Harmonic rejection was achieved with Pt coated mirrors. Slits were opened to provide a beam size at sample of ~3×1 mm^2^ and minimize the photon density, in order to avoid radiation damage. The energy was calibrated over an Ag foil mounted downstream to the sample. XAS spectra were acquired in fluorescence mode, with a 13-element Ge detector from Camberra. Data processing and linear combination analysis were performed with the Athena software;^20^ the chosen fitting range for linear combination analysis was [-20; 80] eV with respect to the edge energy.

### 2.9. STEM-EDX

For the chemical identification of AgNPs, thin sections were observed in scanning/transmission electron microscopy (STEM) mode and analyzed by EDX using either a Zeiss Merlin microscope operated at 30 kV and fitted with two Brucker SDD detectors or a FEI/Tecnai Osiris microscope operated at 80 kV.

### 2.10. FIB-SEM acquisition and data analysis

Focused Ion Beam (FIB) tomography was performed with a Zeiss NVision 40 dual-beam microscope. In a first step, the surface of the sample was protected by the deposition in the microscope of a 2 μm thick Pt or C layer, on a surface of typically 20×30 μm^2^. The resin-embedded HepG2/C3a spheroids were then cut in cross-sections, slice by slice, with a Ga+ ion beam (typically with a current of 300 or 700 pA at a beam acceleration of 30 kV). After a thin slice was removed with the ion beam, the newly exposed surface was imaged in SEM at 1.5 kV using the in-column EsB backscatter detector. For each slice, a thickness of 8 nm was removed, and the SEM images were recorded with a pixel size of 4×4 nm^2^. The image stack was then registered by cross-correlation using the StackReg plugin in the Fiji software.^21^ Manual segmentation was performed in Ilastik^22^ and projected in three-dimension using AVIZO (FEI, USA).

### 2.11. Reflection confocal microscopy

HepG2 cells were seeded on two-well Thermo Scientific^™^ Nunc^TM^ Lab-Tek^TM^ chambered coverglass suitable for confocal microscopy. At 60% of confluency, cells were incubated with 1 μM Hoechst 33342 for 30 minutes at room temperature then washed with complete medium. Confocal microscopy was carried out in a perfusion chamber (POC Chamber, LaCom^®^, Erbach, Germany) and incubation system (O_2_-CO_2_-°C, PeCom^®^, Erbach, Germany) that was mounted on a Leica TCS SP2 AOBS inverted laser scanning confocal microscope using a 63x oil immersion objective (HCX PL APO 63.0 × 1.40 OIL). Cells were exposed to cit-AgNPs or PVP-AgNPs at the indicated concentrations of total Ag just before the first image (t0) then images were acquired on the confocal microscope every 30 minutes and up to 8 hours for cit-AgNPs and 4h for PVP-AgNPs, respectively. For each time-point, a z-stack of 26 pictures with a 1 μm z-step was captured. The laser excitation was set at 351-364 nm for Hoechst, and 488 nm (25%) for the reflection settings. The fluorescence emission was set at 410-485 nm for Hoechst. The detector for reflected light was set at 480-500 nm. For co-localization with Lamp1, after the incubation with AgNPs, cells were fixed with formalin (Sigma) for 30 minutes at room temperature. Fixed cells were permeabilized with 0.5% Triton X-100 in Tris-Buffered Saline containing 0.05% Tween-20 for 60 minutes at room temperature and then blocked with 0.1% Triton X-100 / 3% BSA in PBS containing 0. 05% Tween20 for 30 minutes at room temperature. Blocked cells were incubated with antibody against Lamp1 (Santa-Cruz, sc20011) diluted 1:50 in antibody buffer (0.1% Triton X-100 / 1% BSA / 0.05% Tween20 in PBS) for 60 minutes at room temperature. Cells were then incubated with Alexa Fluor 488 goat anti-mouse secondary antibody diluted 1:1000 (Life Technologies) in antibody buffer for 60 minutes at room temperature. For all the above-mentioned experimental conditions, cells were washed in PBS in between subsequent steps. Cells were finally imaged using Leica TCS SP2 AOBS inverted laser scanning confocal microscope using a 63x oil immersion objective (HCX PL APO 63.0 × 1.40 OIL) as for live imaging.

### 2.12. Statistical analysis

All the quantitative data presented with a standard deviation are based on at least three biological replicates except for XRF experiments that are based on at least three different cells. The statistical comparison of qPCR quantification is based on a standard student t-test. The method of quantification for FXR and LXR activation is described above in this experimental section.

## 3. Results and discussion

### 3.1. Distribution of Ag species within hepatocytes under exposure to low AgNP concentrations

In this study, we used 20 nm citrate-coated AgNPs (cit-AgNPs) and 90 nm PVP-coated AgNPs (PVP-AgNPs) that formed objects of 90 and 160 nm size, respectively, in cell culture media^9^ (Table S1, ESI†). We had previously identified concentrations of 50 and 100 μM as subtoxic in HepG2 cells for citrate- and PVP-coated AgNPs, respectively.^9^ In the current study, the main part of the work was performed at concentrations 4-fold lower, *i.e.* 12.5 μM (cit-AgNP) and 25 μM (PVP-AgNP) that led to similar Ag uptake in 24 hours. These are also non toxic concentrations that did not show significant stress based on heme oxygenase (HMOX) expression (Fig. S1A, ESI†). Indeed, HMOX is regulated by Nrf2, a redox stress sensor, usually activated by metal NPs.^23, 24^ In the chosen conditions, the only marker activated by these low AgNP concentrations was metallothionein (MT) a classical marker of metal ion release into the cytosol (Fig. S1B, ESI†).

In order to visualize Ag in both ionic and particulate forms, we developed a method using X-ray fluorescence (XRF) nano-imaging performed on thin cell sections with the capability to localize these species at the organelle level. The 200 nm sections yielded extremely detailed images of the inside of the cell with the clear identification of various vesicles and compartments by Os staining (Fig. 1A). In HepG2 cells exposed to low concentrations of AgNPs over three days, Ag hot spots corresponding to dissolving NPs were observed in vesicles (Fig. 1B, red-yellow pixels). Using confocal microscopy, AgNPs, both citrate and PVP, can be detected in reflection mode (Fig. S2, ESI†, red) and they were often observed co-localized with Lamp1 stained vesicles, thus identifying them as lysosomes (Fig. S2, ESI†, green, co-localization red-green highlighted with white arrows). In addition, in the same experimental conditions similar perinuclear lysosome vesicles containing dark particles were observed by STEM (Fig. S3-S4, ESI†). EDX showed that they contained Ag as well as sulfur both in the case of cit-AgNPs and PVP-AgNPs (Fig. S3-S4, ESI†) with area of strong co-localization (for instance in Fig. S4B, ESI†, lower panels comparing Ag and S EDX maps). Therefore, AgNPs are transformed within lysosomes. X-ray fluorescence images are consistent with this, since different levels of intensity in Ag are visualized within lysosomes. Moreover, XRF showed that vesicles containing AgNPs had a much higher soluble Ag level compared to the cytosol where it was homogeneously low (Fig. 1B, green compared to light blue pixels). Therefore, Ag(I) species released from AgNPs inside lysosomes are not released freely from these vesicles but are under the control of limiting transport mechanisms that may involve metal-ion transporters such as Cu transporters of the Ctr family.^25^ The cell nucleus was easily identified by its low retention of Os staining (Fig. 1A, noted n) and a low but significant signal for Ag was observed in this organelle (Fig. 1B). In order to perform an accurate elemental quantification, 400 nm thick sections were used. Ag was observed in the nuclei of cells exposed for 48, 72 and 96 hours to cit-AgNPs (Fig. 1C-D and S5A-B, ESI†) and 72 hours to PVP-AgNPs (Fig. S5C, ESI†). Moreover, the superimposition of Os (green) and Ag (red) maps revealed areas in the nucleus with higher Ag content (Fig. 1C-D and S5A-C, noted N, ESI†). These regions were putatively identified as nucleoli, which are usually observed as darker areas by electron microscopy.^26^ The presence of Ag in these areas was confirmed by a specific XRF Ag-Kα peak (Fig. S6, ESI†).

**Fig. 1.**
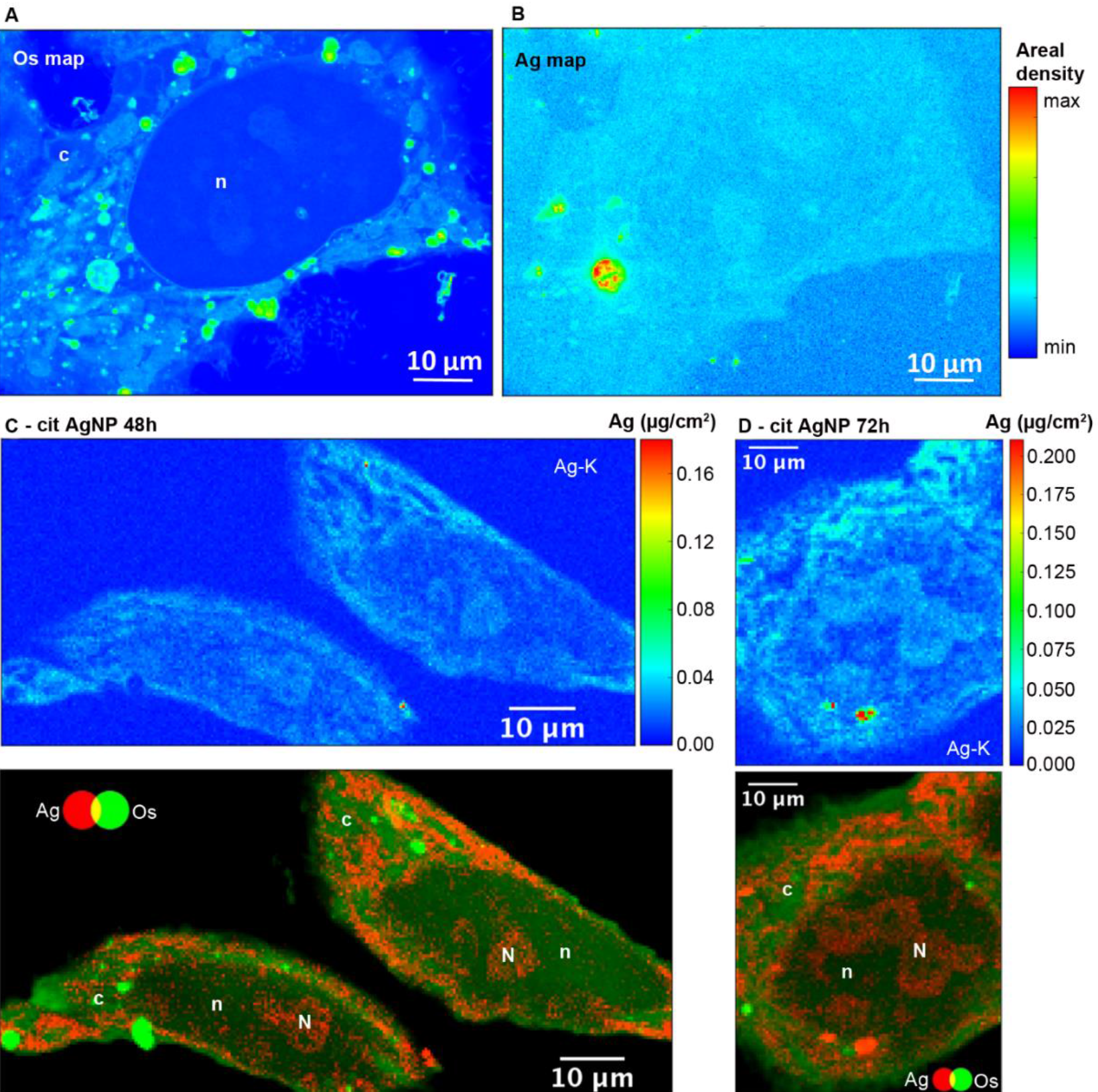
Semi-thin sections of HepG2 cells analyzed by Nano-XRF. (A) Os and (B) Ag areal density maps for a 200 nm section of the same cell exposed for 72 h to 12.5 μM cit-AgNPs. (C) Ag areal density map (upper panel) and two-color map (Ag in red and Os in green, lower panel) of the same region for a 400 nm section of cells exposed for 48 h to 12.5 μM cit-AgNPs. (D) Ag areal density map (upper panel) and two-color map (Ag in red and Os in green, lower panel) of the same region for a 400 nm section of cells exposed for 72 h to 12.5 μM cit-AgNPs. Areal densities are expressed in μg/cm^2^; pixel size is 100×100 nm^2^. Scale bars are 10 μm. *c*: cytoplasm; n: nuclei; N: nucleoli. The upper areal density limit in color bars is set to values that allow the visualization of the low areal density signal.

Between 48 and 72 hours of exposure (Fig. 2A), Ag concentrations increased in the nuclei and in the putative nucleoli structures, while a slight decrease occurred after 72 hours. In the case of cit-AgNP, the amount of Ag per million cells increased at a slower rate than with PVP-AgNPs, and both reached a plateau around 0.7-0.8 μg Ag/10^6^ cells that corresponds to about 4.10^9^ Ag atoms per cell (Fig. S7, ESI†). The uptake of cit-AgNPs between 72 and 96 hours of exposure is thus limited. It is therefore possible that internalized silver is distributed to other organelles between 72 and 96 hours explaining this slight decrease in the nucleus.

**Fig. 2.**
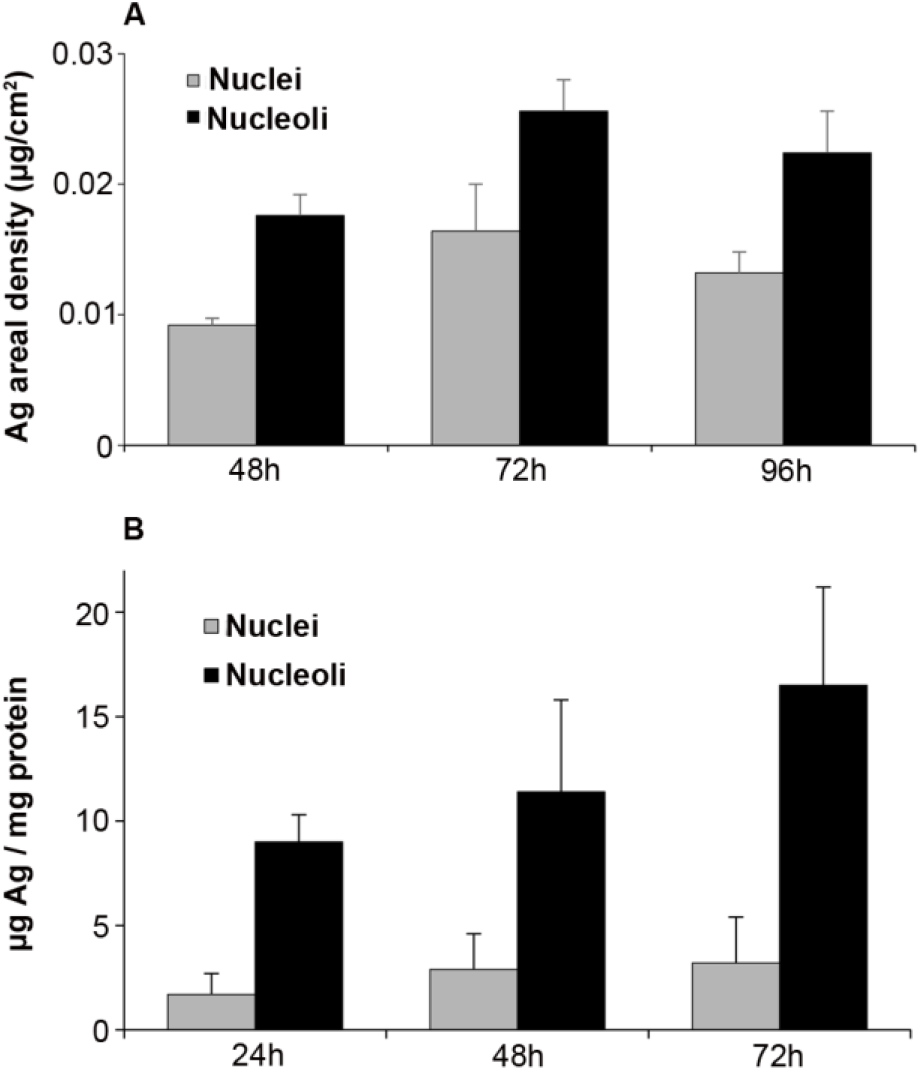
Ag quantification in nuclei and nucleoli. (A) average Ag areal density from Nano-XRF maps in nuclei and nucleoli from HepG2 cells exposed to 12.5 μM cit-AgNPs for 48, 72 or 96 hours. The quantification is based on at least 3 cells. (B) Amount of Ag in nuclei and nucleoli purified from HepG2 cells exposed to 12.5 μM cit-AgNPs for 24, 48 and 72 hours. The quantification was normalized based on the amount of protein and each experiment was repeated at least three times independently.

In order to confirm these data, we quantified Ag in nuclei and bound to nucleoli following subcellular fractionation after 24, 48 and 72 hours of exposure. In order to avoid contamination of the nuclei fraction by endosomes and lysosomes loaded with dense AgNPs, sucrose/percoll density gradient centrifugation was performed. A continuous uptake of Ag in nuclei was observed over 72 hours (Fig. 2B). Nucleoli clearly accumulated Ag since elemental quantification normalized on protein amount showed about five times higher Ag concentrations compared to the total nuclei fraction and with the same kinetic trend (Fig. 2B). This quantification is not absolute and the ratio nucleoli/nuclei is higher than that based on XRF quantification but both methods show consistently a higher amount of Ag in nucleoli.

Altogether, our study is the first to show that Ag(I) species enter nuclei and are preferentially accumulated in nucleoli. These novel results are supported by the identification of proteins from the nucleoli possessing Ag-affinity *in vitro,* the so-called AgNOR (Nucleolar Organizer Region)^27, 28^ that most probably bind Ag *via* stretches of glutamate or aspartate residues.^29^ However, carboxylates do not possess a high affinity for Ag(I), making Ag-rich nucleoli a reservoir of Ag(I) that can be eventually released in the nucleus and bind higher affinity protein sites.

### 3.2. Impact of Ag(I) species on nuclear functions

The initial phase of ribosome biogenesis takes place in nucleoli,^28, 30^ in particular with the biosynthesis of the 45S pre-ribosomal RNA (pre-rRNA). We assessed a possible impact of Ag binding to nucleolar proteins on this function. The amount of 45S pre-rRNA was determined by quantitative real-time PCR (qPCR) in HepG2 cells exposed or not to cit-AgNPs at 6.25 and 12.5 μM for up to 4 days (Fig. 3A). 45S pre-rRNA production remained almost constant in the tested conditions. We only observed a slight overexpression after 24 hours of exposure to cit-AgNPs at 12.5 μM. It is possible that Ag binding to AgNOR slightly disturbs nucleolar functions but as ribosomes are essential, cells have no doubt developed ways to efficiently correct for alteration in their biogenesis pathway. In addition, Ag binding to AgNOR occurs in domains with unknown functions for these proteins, which could explain a limited impact on ribosome biogenesis.

**Fig. 3.**
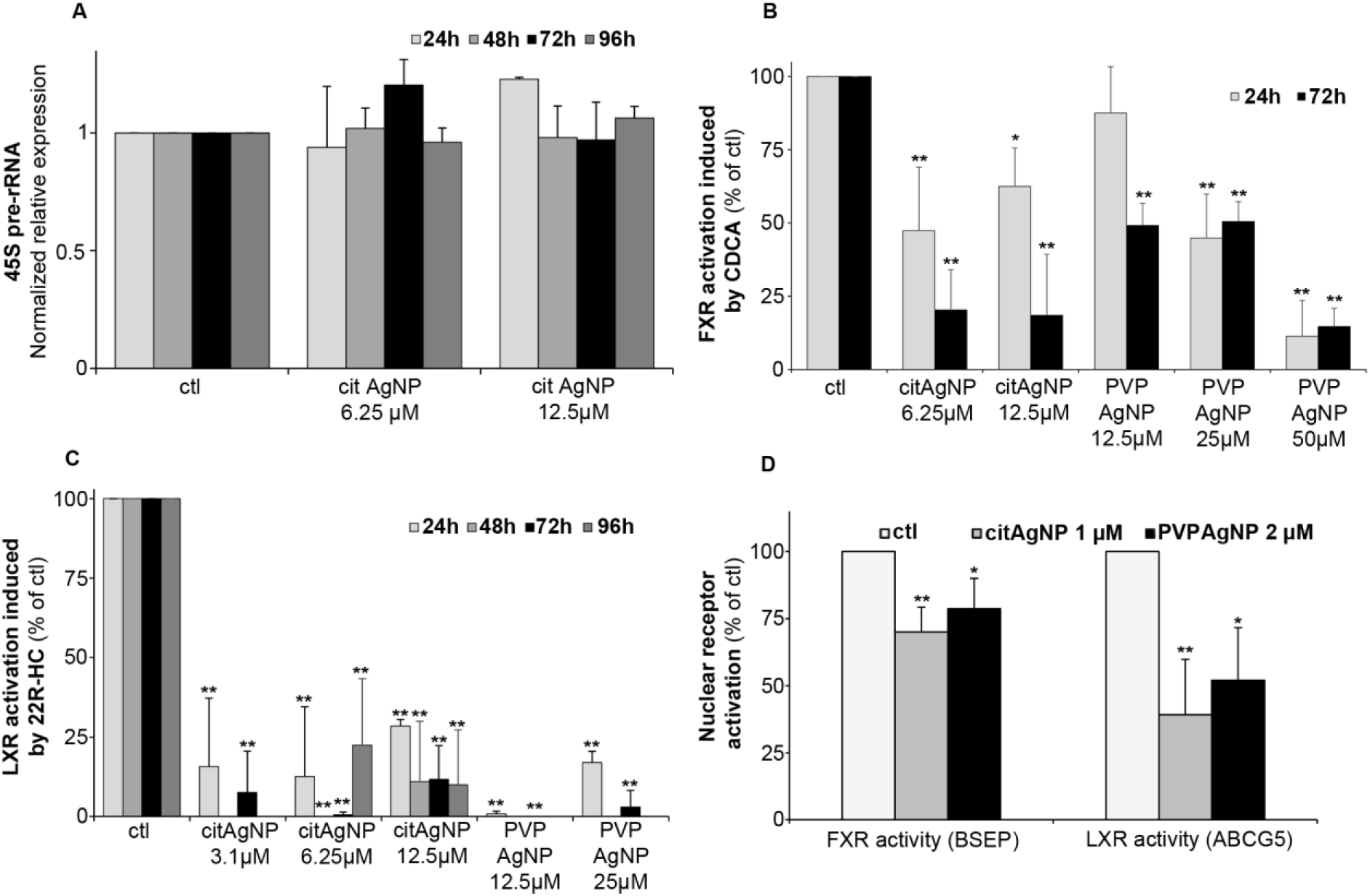
Impact of AgNP exposure on 45S pre-rRNA expression and FXR and LXR activity. (A) variation of 45S pre-rRNA expression levels in HepG2 cells exposed to 6.25 or 12.5 μM cit-AgNPs for 24, 48, 72 or 96 hours. The amount of RNA was determined by qPCR with unexposed cells (ctl) set at 1. (B) FXR activity followed by CDCA-induced BSEP expression. HepG2 cells were exposed or not to AgNPs for 24 or 72 hours and to CDCA at 80 μM or DMSO during the last 17 hours. The RNA levels of BSEP were quantified in the different conditions using qPCR. 100% FXR activity was set using control conditions (unexposed to AgNPs) by calculating the ratio: amount of BSEP-RNA in ctl+CDCA / amount of BSEP-RNA in ctl without CDCA. This ratio (+CDCA/-CDCA) was measured for each condition and divided by the ctl ratio in order to determine the percentage of FXR activity in cells exposed to AgNPs (for details see experimental section (C) LXR activity followed by 22R-HC-induced ABCG5 expression. HepG2 cells were exposed or not to AgNPs for 24, 48, 72 or 96 hours and to 22R-HC at 10 μM or ethanol during the last 20 hours. The RNA levels of ABCG5 were quantified in the different conditions using qPCR. 100% LXR activity was set using control conditions (unexposed to AgNPs) by calculating the ratio: amount of ABCG5-RNA in ctl+22R-HC / amount of ABCG5-RNA in ctl without 22R-HC. This ratio (+22R-HC/-22R-HC) was measured for each condition and divided by the ctl ratio in order to determine the percentage of LXR activity in cells exposed to AgNPs (for details see experimental section). (D) FXR and LXR activities measured as in B and C in HepG2 cells exposed or not for 10 days to 1 μM cit-AgNPs or 2 μM PVP-AgNPs. For A to D each experiment was performed at least three times independently. * stands for data statistically different from the corresponding control with p<0.05 and ** stands for data statistically different from the corresponding control with p<0.01.

The nucleus carries out many critical functions and any disruption can play a role in the development of non-transmissible pathologies such as cancers. Environmental pollutants, those classified as endocrine disruptors for instance, impair the activity of some nuclear receptors (for review see^31^). These are ligand-activated transcription factors with finely tuned activity. They contain zinc finger DNA binding domains that are structured by zinc ions coordinated by four cysteines (Cys). This amino-acid possesses thiol functions that are very good ligands for Cu(I) or Ag(I) ions and it has been observed *in vitro* that Cu(I) can displace Zn(II) from zinc finger domains.^32–34^ Recently, it was shown that an excess of intracellular Cu(I) in Wilson disease impairs the function of nuclear receptors in the liver.^35–37^ To assess the activity of crucial liver nuclear receptors, we determined transcript levels of target genes under the control of these receptors upon addition of their specific ligands in standard growth conditions (100% activation) versus those in cells exposed to AgNPs (Fig. 3B-D). Farnesyl X receptor (FXR) is critical in maintaining bile salt homeostasis, and the addition of chenodeoxycholic acid (CDCA) in the culture media triggers the expression of bile salt export protein (BSEP) mRNA. The exposure of HepG2 cells to cit-AgNPs at 6.25 or 12.5 μM for 24 hours inhibited about half of FXR’s activation and a prolonged exposure to 72 hours led to 80% inhibition (Fig. 3B). PVP-AgNPs possessed a lower inhibitory capacity leading to only about 50% inhibition after 72 hours at 12.5 and 25 μM, and 50 μM were required to inhibit more than 85% of FXR’s activity (Fig. 3B). The level of FXR mRNA expression was also determined by qPCR in the different exposure conditions (Fig. S8A, ESI†). It was observed that CDCA itself induced a slight down-regulation of FXR expression most probably due to a feedback loop avoiding hyper-activation of FXR. The CDCA-induced down-regulation was observed in all tested conditions except at 50 μM PVP-AgNP. Moreover, cit-AgNPs at 12.5 μM and PVP-AgNPs at 50 μM induced a moderate and high down-regulation of FXR, respectively, most probably because of the stress induced to hepatocytes by this concentration of AgNP exposure as shown by HMOX low and moderate mRNA overexpression, respectively (Fig. S1A, ESI†). Overall, for an unknown reason FXR is more sensitive to cit-AgNPs than to PVP-AgNPs.

Liver X receptor (LXR) is a key nuclear receptor for liver metabolism as it is a major regulator of glucose, lipid and cholesterol metabolisms. LXR can be activated by various ligands including cholesterol derivatives such as 22R-hydroxycholesterol (22R-HC) that induces the expression of ABCG5, an ABC transporter involved in biliary cholesterol excretion. Just like FXR, the exposure of HepG2 cells to cit-AgNPs for only 24 hours at 6.25 or 12.5 μM inhibited 70-85% of LXR’s activity (Fig. 3C). Moreover, a 48-hour exposure to AgNPs was enough to completely abolish LXR activity (Fig. 3C) and a slightly lower inhibition was observed for a 96 hour exposure to 6.25 μM, most probably because of the redistribution of Ag throughout the cell as shown by Nano-XRF (Fig. 2A). Similar results were observed with PVP-AgNPs with a complete inhibition of LXR at 12.5μM (Fig. 3C). In order to exclude a direct effect on LXR expression, we quantified LXR mRNA by qPCR. For LXR, we were able to detect both LXRα and LXRβ mRNAs in HepG2 cells and we choose to follow LXRβ expression in our various exposure conditions (Fig. S8B, ESI†). In contrast to FXR, the ligand 22R-HC did not induce a down-regulation of LXRβ expression. Cit-AgNPs at 3.1 μM did not affect LXRβ expression, while concentrations of 6.25 and 12.5 μM induced only a slight down-regulation between 24 and 72 hours of exposure that was lost for the 96-hour exposure (Fig. S8B, ESI†). In the case of PVP-AgNP exposure, only 12.5 μM for 72 hours induced a slight down-regulation of LXRβ, while the level of mRNA was quite stable in all other conditions.

The concentration of cit-AgNPs was then decreased down to 3.1 μM of Ag in order to observe effects on LXR activity that could not be related to metal stress in hepatocytes. In these conditions, MTs were only minimally overexpressed (Fig. S1B, ESI†) and LXRβ was very stable (Fig. S8B, ESI†). Interestingly, such a low concentration of AgNPs led to about 80 and 90% inhibition of LXR after 24 and 72 hours of exposure (Fig. 3C), respectively. Thus, we prove that the exposure to low and non-toxic concentrations of AgNPs impairs nuclear receptor activity in hepatocytes with an extent of inhibition that is receptor-dependent. Indeed, we observed a complete inhibition of LXR and only a partial inhibition of FXR that could be due to different subcellular localizations of these two nuclear receptors in basal conditions.

Finally, in order to very closely mimic chronic exposure to low but constant levels of Ag, we studied exposures to 1 μM cit-AgNPs or 2 μM PVP-AgNPs that led to a low but detectable intracellular Ag concentration. Its level stabilized after 7 to 10 days around 45 and 70 ng Ag/10^6^ cells (between 2.5 and 4.10^8^ Ag atoms per cell) for cit-AgNPs and PVP-AgNPs, respectively (Fig. S9, ESI†), while lower and higher concentrations delivered undetectable and time-dependent increasing amounts of Ag in hepatocytes, respectively. In the latter case, critical Ag levels were reached at one point making it impossible to work in chronic-mimicking conditions as AgNP exposure concentrations ended up being too high, while 1 μM cit-AgNPs and 2 μM PVP-AgNPs were ideal to approach this objective. Similar effects were observed with 1 μM cit-AgNP and 2 μM PVP-AgNP exposure with only a moderate inhibition of FXR, 30 and 20%, respectively, and a significant inhibition of LXR, 60 and 50%, respectively (Fig. 3D). These results confirmed the lower sensitivity of FXR to Ag(I) and also proves that chronic exposure to AgNPs delivered sufficient amounts of Ag(I) to disrupt nuclear receptor activity. Overall, the experiments on nuclear functions did not show major differences between AgNPs with different coatings except a slightly higher concentration required for PVP to reach the same level of toxicity.

### 3.3. From AgNP endocytosis to Ag(I) nuclear entry, a rapid process

Our data have clearly demonstrated the rapid uptake by nuclei of a significant amount of Ag(I) ions that accumulate in nucleoli and induce dose-dependent effects on nuclear functions (Fig. 1–3). A debate is still ongoing concerning the possible uptake of the nanoparticulate form of AgNPs in the nuclei, but the methods used to demonstrate this hypothesis cannot be considered as conclusive (for detailed explanations, see the recent review ‘). In order to deepen our understanding of AgNP trafficking towards nuclei, we performed a combined confocal reflected light microscopy and 3D-EM analysis using FIB-SEM. The particle nature of AgNPs makes it possible to visualize their dynamics by reflectance confocal microscopy (Fig. 4 and Fig. S10-S12, ESI†). Cit-AgNPs could only be detected in cells three hours after exposure, while PVP-AgNPs were observed as clear single spots immediately following their addition (Fig. 4A and S10, ESI† *versus* 4B and S11, ESI†). This is consistent with our previous observations of isolated PVP-AgNPs as opposed to agglomerated cit-AgNPs during endocytosis^9^ and with the kinetics of uptake of both types of AgNPs (Fig. S7, ESI†). We also recently published electron microscopy micrographs revealing PVP-AgNPs bound to the cell membrane at the interface between cells.^1^ This observation was made on thin cell sections and was specific to PVP-AgNPs. Similarly, confocal reflectance microscopy images showed particles evenly distributed on lines or curves between cells (Fig. 4B and S11, red arrows) in addition to intracellular particles. Therefore, for an unknown reason, we confirmed that a fraction of the PVP-AgNPs but not of the cit-AgNPs bind to membranes between cells as described in Marchioni *et al*.^1^ For cit-AgNPs, the reflection signal in the cell increased over time with a tendency to accumulate in the perinuclear region (Fig. 4A and S10, white arrows, ESI†). In the case of PVP-AgNPs, the dynamics were faster and NPs that came in the vicinity of nuclei disappeared rapidly (Fig. 4B and S11, white arrows, ESI†). Therefore, within just a few hours, AgNPs reached the vicinity of the nucleus where they promptly dissolved. In order to go a step further, PVP-AgNP distribution within hepatocytes grown as spheroids was also obtained at nanometer resolution using FIB-SEM. This method provided the unique capability to observe individual NPs without any preparation-dependent artifacts that might lead to erroneous localization of NPs within the cell. Indeed, the biological sample is embedded in a resin and imaged 8 nm layer by 8 nm layer thanks to Ga gun nanomachining (for review see^38^) thus avoiding diamond knife alterations of AgNP location. Moreover, for this experiment, a derivative of the HepG2 cell line, named HepG2/C3a,^39^ was used to grow hepatocytes in 3D as spheroids as this cell line is capable of forming functional bile canaliculi in 3D culture, providing a model closer to functional tissue. Similarly to our data on 2D cell culture, FXR activity was readily inhibited in hepatocytes grown as spheroids after 48 hours of exposure to 12.5 μM of cit-AgNPs or 25 μM of PVP-AgNPs (Fig. S13, ESI†) as observed in 2D culture (Fig. 3B). These results increase the relevance of our data on nuclear receptor inhibition upon AgNP exposure. FIB-SEM analysis revealed that PVP-AgNPs were only observed in vesicles, most probably endosomes and/or lysosomes (Fig. 5A white arrows and Video S1, ESI†). Moreover, in accordance with our previous data (Fig. S3-S4, ESI†), AgNPs were frequently found in lysosomes close to the nucleus, as shown in the 3D reconstruction of a typical hepatocyte (Fig. 5B-C and Video S1, ESI†) as well as in thin sections analyzed by STEM-EDX that confirmed the dark particles to be transformed AgNPs, since Ag and S were detected (Fig. S14, ESI†). These results confirmed reflection confocal microscopy data adding nanometer resolution. Thorough analysis of the original SEM micrographs used to make the 3D reconstruction confirmed the proximity between AgNP-containing vesicles and the nucleus, showing close contacts between both membranes (Fig. 5D, red arrow). However, AgNPs were never observed inside nuclei.

**Fig. 4.**
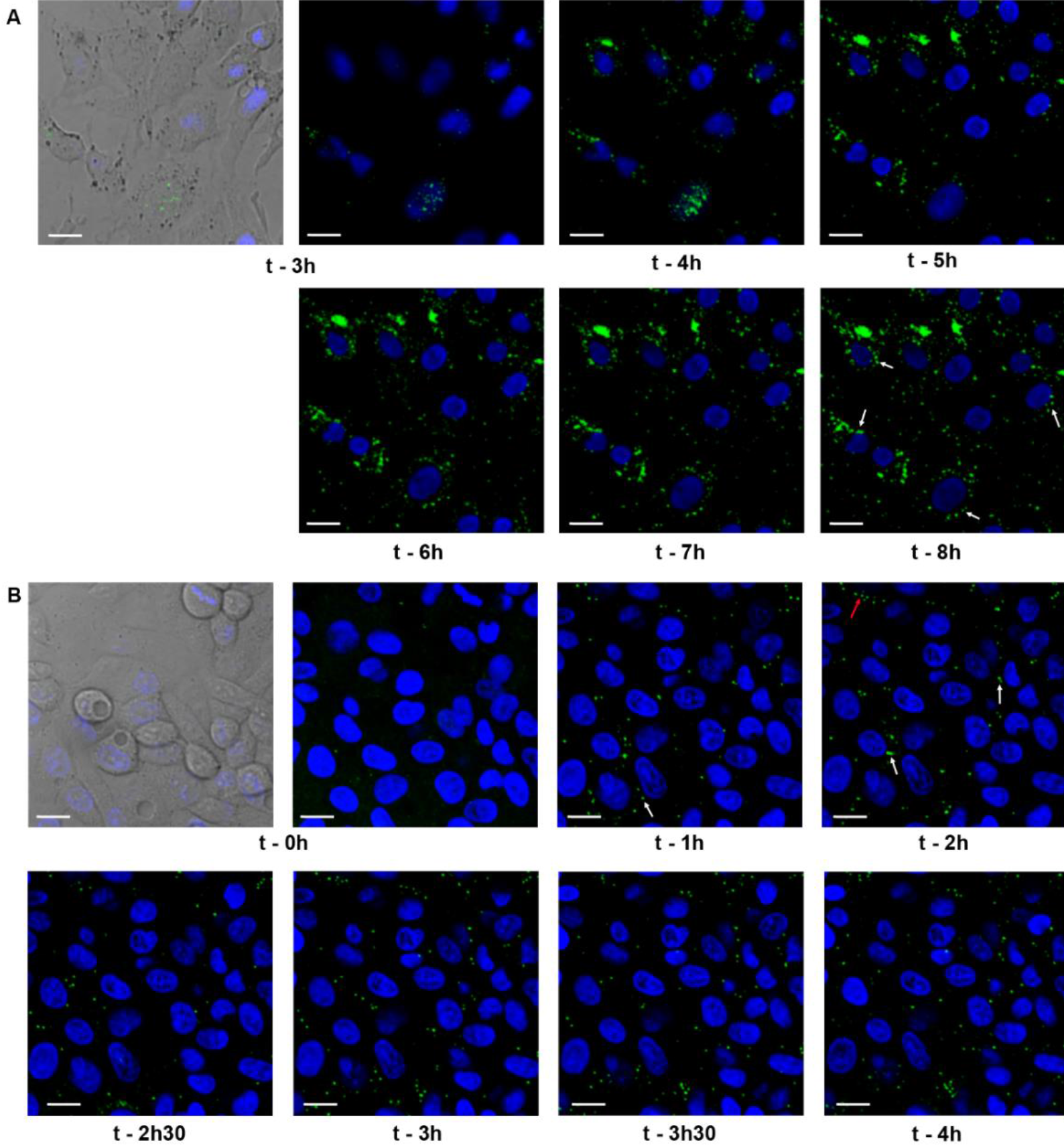
Dynamics of cit-AgNPs and PVP-AgNPs in HepG2 cells. (A) confocal fluorescence and reflectance microscopy of HepG2 cells exposed to 12.5 μM of cit-AgNPs for the indicated times. Nuclei were labeled with Hoechst (blue) and AgNPs were observed in reflection mode (green). At t=3h, a bright field image is also shown. (B) confocal fluorescence and reflectance microscopy of HepG2 cells exposed to 25 μM of PVP-AgNPs for the indicated times. Particles are represented in green and nuclei in blue. At t=0h, a bright field image is also shown. White arrows pinpoint particles close to the nucleus and red arrows PVP-AgNPs bound to the cell membrane. The scale bars correspond to 20 μm.

In addition, the proportion of Ag(I) and AgNPs in hepatocytes was analyzed using X-ray absorption spectroscopy using the spectra of a AgNP suspension and of a Ag-GSH complex as standards (Fig. S15, ESI†). The spectra of cells showed similar features, suggesting similar dissolution fractions under the explored exposure conditions. The main features of the AgNP reference spectrum were present in the spectra of exposed cells, but heavily smoothed as a consequence of the predominance of Ag-thiolate species (represented by the Ag-GSH reference compound), as observed previously in identical experimental conditions.^8, 9^ The semi-quantitative evaluation of the different Ag species in cells obtained by linear combination fitting (Table S3, Fig. S15 red curves, ESI†) confirmed an extensive dissolution for both cit-AgNPs (12.5 μM) and PVP-AgNPs (25 μM) after only 24 hours of exposure leading to 70 and 76% of Ag(I)-thiolate complexes, respectively. This dissolution process was maintained over time because 77% of uptaken cit-AgNPs were dissolved after 48 hours of exposure. It is therefore interesting to notice that lower exposure concentrations lead to higher dissolution of AgNPs into Ag(I) ions since we previously showed that no more than 50% of internalized AgNPs were dissolved after 24h exposure to 100 μM AgNPs.^9^ Altogether, these data showed that AgNPs rapidly dissolved upon uptake and continuously over time under exposure to low and non-toxic concentration of AgNPs.

This extent of dissolution is consistent with the observed perinuclear localization of AgNP-containing endolysosomes. Indeed, the closer lysosomes are to the nucleus, the lower their internal pH^40^ and lower pH values favor AgNP dissolution. Nonetheless, the combination of all our data prompted us to reconsider our vision on AgNP dissolution. Indeed, different forms of species containing Ag and S exist within the cell. XRF microscopy showed intense Ag signal within endosomes and lysosomes and low Ag signal in cytosol and other organelle such as nuclei. However, STEM-EDX experiments revealed that lysosomes contain both AgNPs close to pristine form as well as highly transformed AgNPs (Fig. 5 and Fig. S3-S4, ESI†). This is particularly clear in Fig. 5E-G where darker particles have a high signal in Ag (pink square in Fig. 5F), while grey particle-like areas have a low Ag signal with the presence of S in a similar intensity (blue and green square in Fig. 5F). Interestingly, in Fig. 5F, we can almost see a connection between a lysosome and the nucleus (red arrow) but no particle nor Ag signal was observed in the nucleus even in areas that look darker in STEM. Similarly, Ag was never detected in the cytosol by EDX, while_it was by XRF both in cytosol and nuclei. Species containing Ag and S thus exist in two forms, particle-like in lysosomes and soluble elsewhere. Therefore, our results constitute the first evidence of a rapid trafficking of endosomes maturing into lysosomes that contain AgNPs in conditions favoring their rapid transformation into Ag(I) species, which are then partly transferred as soluble complexes into the nucleus where they bind and inhibit nuclear receptors. Mechanisms governing the shuttling of nuclear receptors between cytoplasm and nucleus are complex and not fully understood,^41, 42^ and are partly receptor-specific. In the case of LXR, the metabolic status of the cell *(i.e.* glucose level) can favor LXR nuclear translocation.^43^ Therefore, the fast kinetics of Ag(I) nuclear transfer provides one possible explanation for the difference in sensitivity to AgNPs between FXR and LXR. Indeed, Ag(I) directly delivered to the nucleus could rapidly and efficiently bind and inhibit nuclear LXR.

**Fig. 5.**
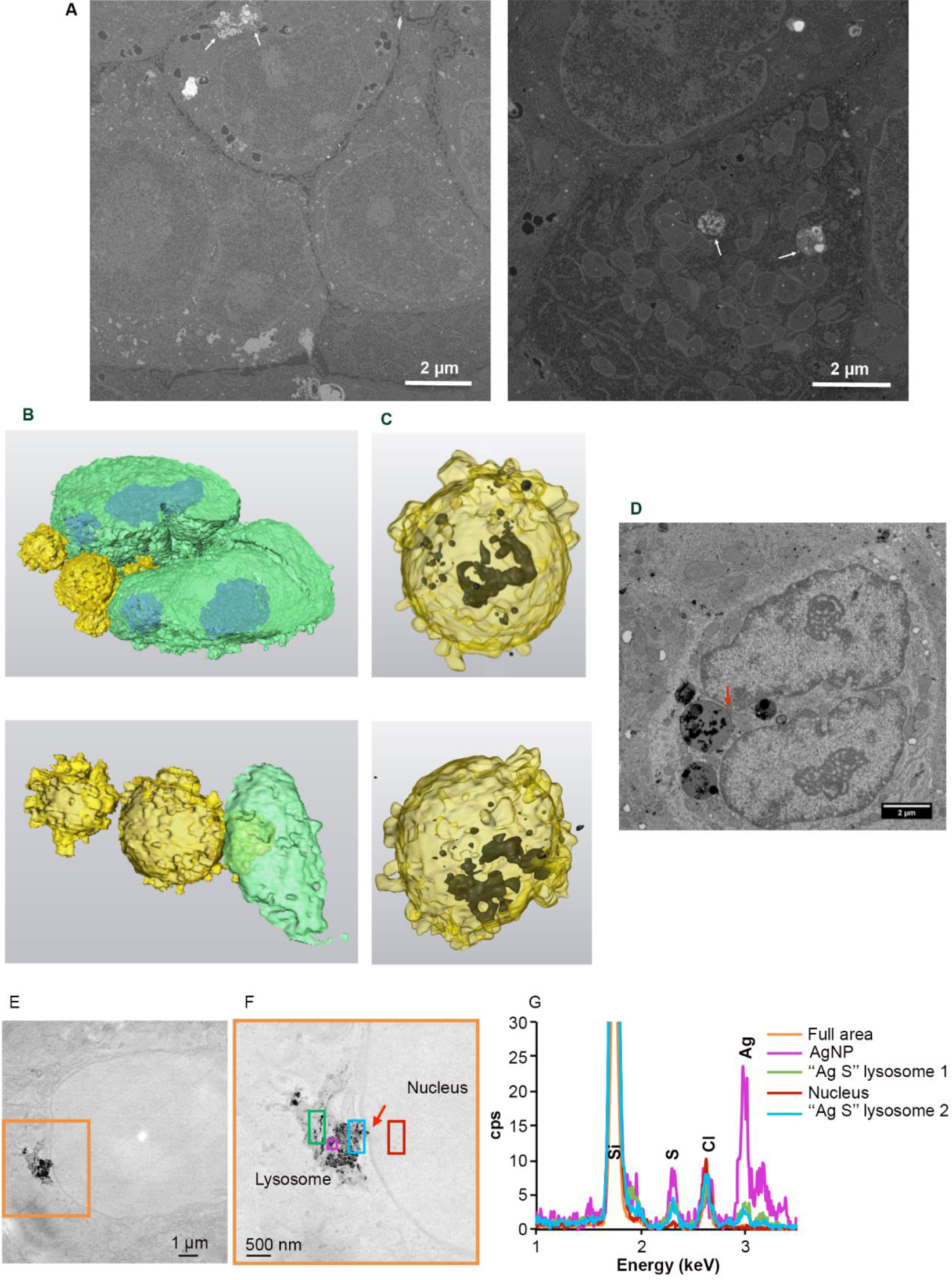
EM analysis of subcellular distribution of AgNP-containing vesicles. (A) Original micrographs extracted from two different FIB-SEM stack of images (see also Video S1 for stack 1 with the corresponding segmentation). Endo-lysosomes containing AgNPs close to the nucleus (left) and in the middle of the mitochondrial network (right) are indicated with white arrows. (B) 3D reconstruction of a nucleus (green) with bound vesicles (yellow) containing AgNPs (black) seen by transparency (C). (D) SEM micrograph from the stack used to make the 3D reconstruction (B, C). This micrograph shows a direct contact between the AgNP-containing vesicle and the nucleus (red arrow). Scale bars represent 2 μm. (E) Large field of view STEM micrograph showing the presence of a lysosome containing NPs in contact with a nucleus in HepG2 cells exposed for 72 hours to 12.5 μM cit-AgNPs. (F) STEM micrograph of the orange area in (E) at higher magnification. The red arrow pinpoint a region of close contact between the lysosome and the nuclei. The different colored boxes correspond to area scanned by EDX with the spectra presented in (G). (G) EDX spectra of different regions in the energy range 1 – 3.5 keV. The position of Si, S, Cl and Ag peaks is highlighted.

## 4. Conclusions

In the current study, we developed an innovative elemental imaging methodology enabling unprecedented localization of both AgNPs and Ag(I) species at the organelle level. AgNPs were only observed in endosomal and lysosomal vesicles, which move towards a perinuclear localization. A combination of imaging methods revealed the transformation of AgNPs into dense species containing Ag and S within lysosomes. Highly sensitive elemental imaging revealed a direct and rapid entry of soluble Ag(I) species into the nucleus where it accumulates, weakly bound to nucleoli, and impairs nuclear receptor activity. Indeed, one day of exposure is sufficient to partially or totally inhibit a set of nuclear receptors. Besides, extremely low doses leading to less than 50 fg of silver per cell is sufficient to inhibit more than half of LXR activity, a key regulator of liver metabolism.

Based on previously published *in vitro* data,^32–34^ we can ascribe the observed effects on nuclear receptor activity to a displacement of Zn(II) by Ag(I) in the zinc finger DNA binding domains, as already proposed with Cu(I) in a mouse model of Wilson disease.^35–37^ Zinc fingers require a tetra-coordination geometry to be able to bind their specific DNA-binding domains upon ligand binding, while Ag(I) would only coordinate two or three Cys leading to an inactive transcription factor. Indeed, it was shown very recently that Ag can interfere with Zn fingers *in vitro*^44^ supporting our results. The impairment of this activity is highly detrimental for liver metabolism as it will affect lipid, cholesterol and bile salt homeostasis, among others. Moreover, released Ag(I) ions could also interact with other nuclear receptors, possibly leading to developmental diseases^31^ such as those observed with endocrine disruptors. It could therefore be of high relevance to study the impact of the combined exposure to AgNPs and endocrine disruptors that target the same protein family. In addition, the binding of released Ag(I) cannot be limited to nuclear receptors. Zinc finger proteins account for 3% of the human genome and many sites with two or three exposed thiols exist in proteins. It would therefore be of high interest to determine the whole Ag metalloproteome upon realistic exposure to dietary or medical AgNPs.

In conclusion, the current work is a pioneer study in understanding physiological disruptions induced by exposure to very low doses of AgNPs mimicking chronic exposure. Further work is now required to precisely determine our exposure levels in order to clearly define an AgNP safety framework. In this context, we recently developed robust, versatile and safer biocides based on assemblies of AgNPs.^45^

Moreover, the idea of using AgNPs in cancer therapy recently emerged and this work on a detailed analysis of AgNP trafficking and transformation in cells can be highly valuable for these types of development. Once again, deciphering toxicity mechanisms of nanomaterials could lead to nanomedecine applications as described recently by Peng *et al*. that made use of nanomaterial-induced endothelial leakiness to favor cancer therapy.^46^

## Acknowledgements

The authors acknowledge the European Synchrotron Radiation Facility for providing beam time (inhouse research on ID16B-NA and experiment LS-2639 on BM23), and O. Mathon for the excellent support during XAS data acquisition. This work used the platforms of the Grenoble Instruct-ERIC Center (ISBG: UMS 3518 CNRS-CEA-UGA-EMBL) with support from the French Infrastructure for Integrated Structural Biology (FRISBI; ANR-10-INSB-05-02) and GRAL, a project of the University Grenoble Alpes graduate school (Ecoles Universitaires de Recherche) CBH-EUR-GS (ANR-17-EURE-0003) within the Grenoble Partnership for Structural Biology (PSB). The IBS Electron Microscopy facility is headed by Guy Schoehn and supported by the Auvergne Rhône-Alpes Region, the Fondation Recherche Medicale (FRM), the fonds FEDER and the GIS-Infrastrutures en Biologie Sante et Agronomie (IBISA). This research is part of the LabEx SERENADE (grant ANR-11-LABX-0064) and the LabEx ARCANE and CBH-EUR-GS (grant ANR-17-EURE-0003). This work was supported by the CEA Transversal Programs Toxicology and Nanoscience through the NanoTox-RX grant and the CEA DRF-Impulsion grant FIB-Bio. This work was also funded by the Université Grenoble Alpes – AGIR grant NanoSilverSafe and IRS NanoBIS grant and by the RHU program ANR-16-RHUS-0005. EK was supported by a CMIRA Accueil Sup 2016 fellowship.

