## Supplementary information for "Visualization of silver nanoparticle intracellular trafficking revealed nuclear translocation of silver ions leading to nuclear receptor impairment"

a. ESRF, The European Synchrotron. 71 avenue des Martyrs, 38000 Grenoble, France

b. Univ. Grenoble Alpes, CNRS, CEA, IRIG, Laboratoire de Chimie et Biologie des Métaux BIG-LCBM, F-38000 Grenoble, France

c. Univ. Grenoble Alpes, CEA, INAC-IRIG, MEM, F-38000 Grenoble, France

d. Institut de Biologie Structurale, CEA, CNRS, Univ. Grenoble Alpes, CEA, CNRS, IBS, 71 avenue des Martyrs, F-38042 Grenoble, France

e. Univ. Grenoble Alpes, INSERM U1055, Laboratory of Fundamental and Applied Bioenergetics (LBFA), and Environmental and System Biology (BEeSy),, 38000 Grenoble, France

f. Integrated Structural Biology Grenoble (ISBG) CNRS, CEA, Université Grenoble Alpes, EMBL, 71 avenue des Martyrs, F-38042, Grenoble, France

g. Institute of Biochemistry and Biophysics, Polish Academy of Sciences, Pawinskiego 5a, 02-106 Warsaw, Poland

h. DRF/BIG-INSERM U1160, Hôpital St Louis, F-75010 Paris

### Supplementary figures

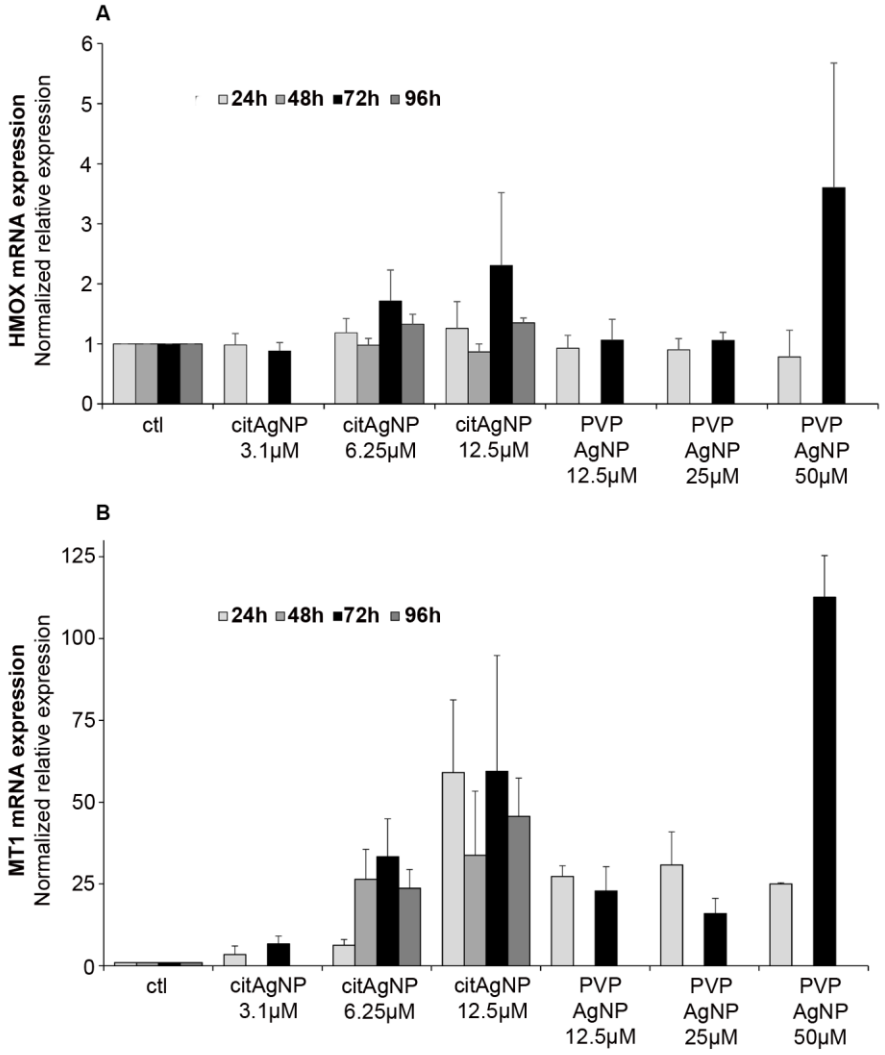


**Fig. S1.** mRNA levels of redox and metal stress indicators. (A) HMOX mRNA fold increase in HepG2 cells exposed to AgNPs. (B) MT1 mRNA fold increase in HepG2 cells exposed to AgNPs. Results are expressed as the relative change in expression compared to control (ctl), corresponding to cells unexposed to AgNPs. The results are expressed as means +/- standard error of the mean of at least three independent experiments.

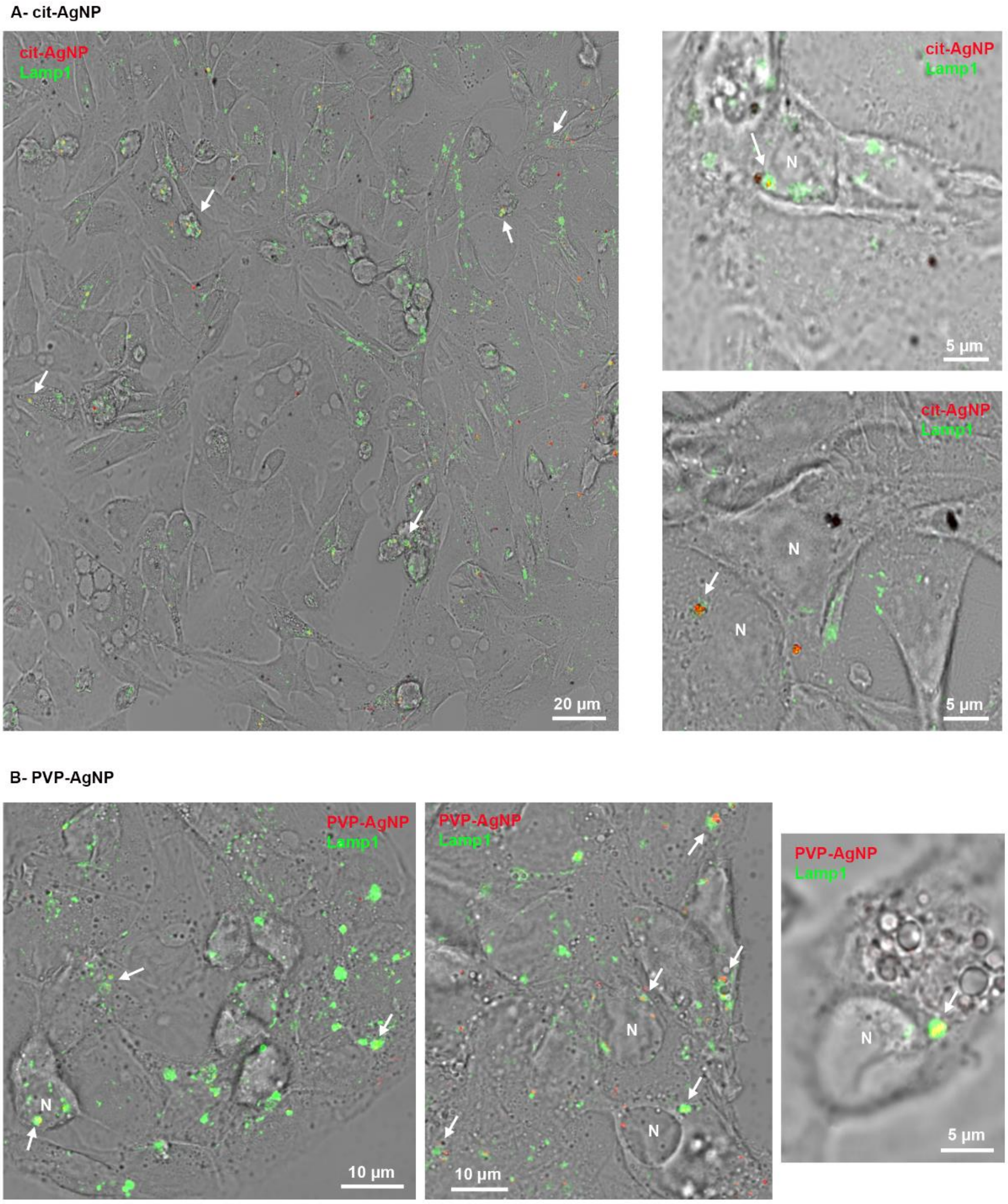


**Fig. S2.** Dynamics of AgNPs in HepG2 cells. Confocal fluorescence and reflectance microscopy of HepG2 cells exposed to 12.5 µM of cit-AgNPs (A) or 25 µM PVP-AgNPs (B) for 24 hours. AgNPs were observed in reflection mode (red) and Lamp1 by immunofluorescence (green). White arrows pinpoint some of the lysosomes (Lamp1) containing AgNPs. N stands for nuclei.

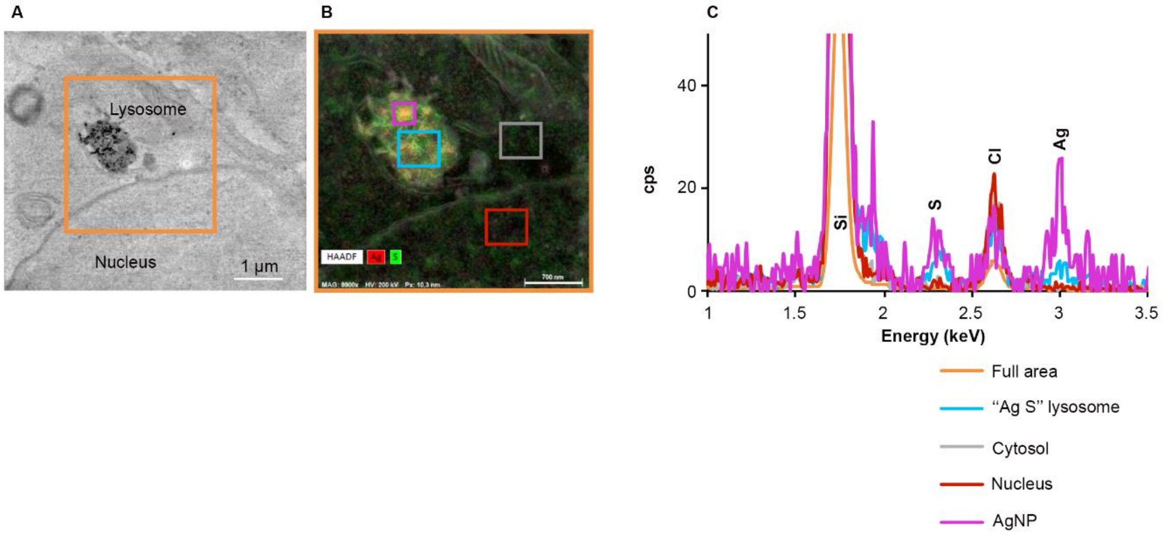


**Fig. S3.** Visualization and analysis of cit-AgNPs in the perinuclear region. (A) Large field of view STEM micrograph showing the presence of a lysosome containing NPs in contact with a nucleus in HepG2 cells exposed for 24 hours to 12.5 µM cit-AgNPs. (B) EDX map of the orange area in (A) showing HAADF map in black and white, Ag map in red and S map in green. The different colored boxes correspond to the area selected for spectra extraction presented in (C). (C) EDX spectra of different regions in the energy range 1 – 3.5 keV. The position of Si, S, Cl and Ag peaks is highlighted.

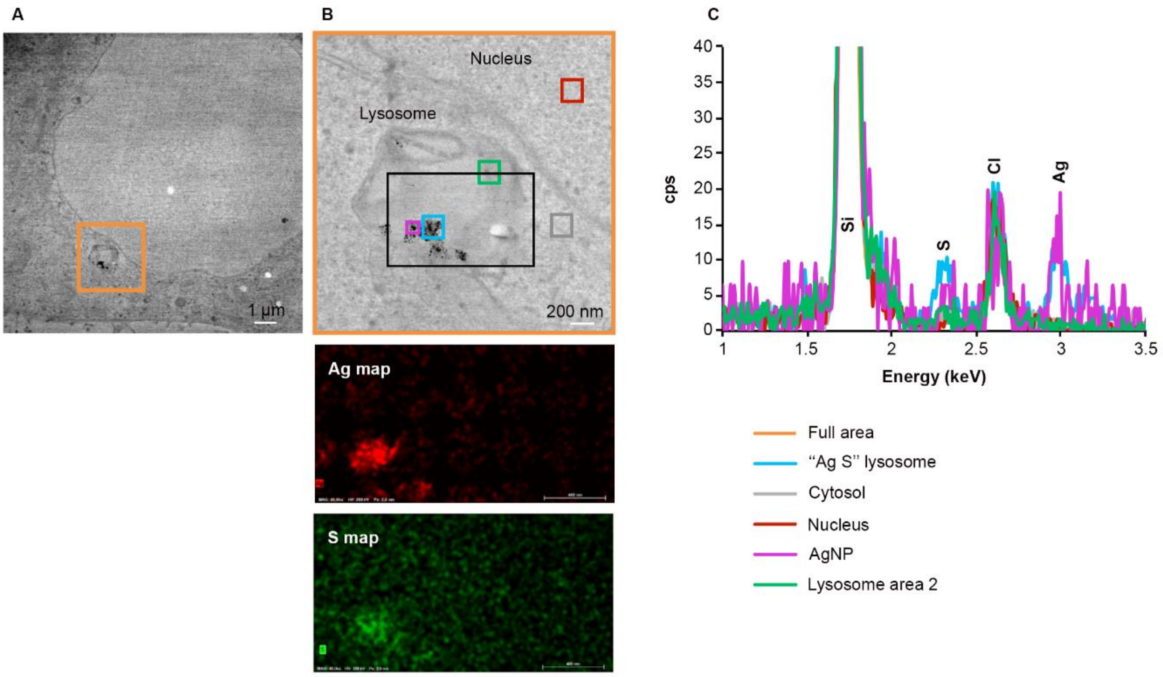


**Fig. S4.** Visualization and analysis of PVP-AgNPs in the perinuclear region. (A) Large field of view STEM micrograph showing the presence of a lysosome containing NPs in contact with a nucleus in HepG2 cells exposed for 24 hours to 25 µM PVP-AgNPs. (B) STEM micrograph of orange area in (A) at higher magnification. The different colored boxes correspond to the area scanned by EDX with spectra presented in (C). Lower panels show EDX Ag and S maps of the black area. (C) EDX spectra of different regions in the energy range 1 – 3.5 keV. The position of Si, S, Cl and Ag peaks is highlighted.

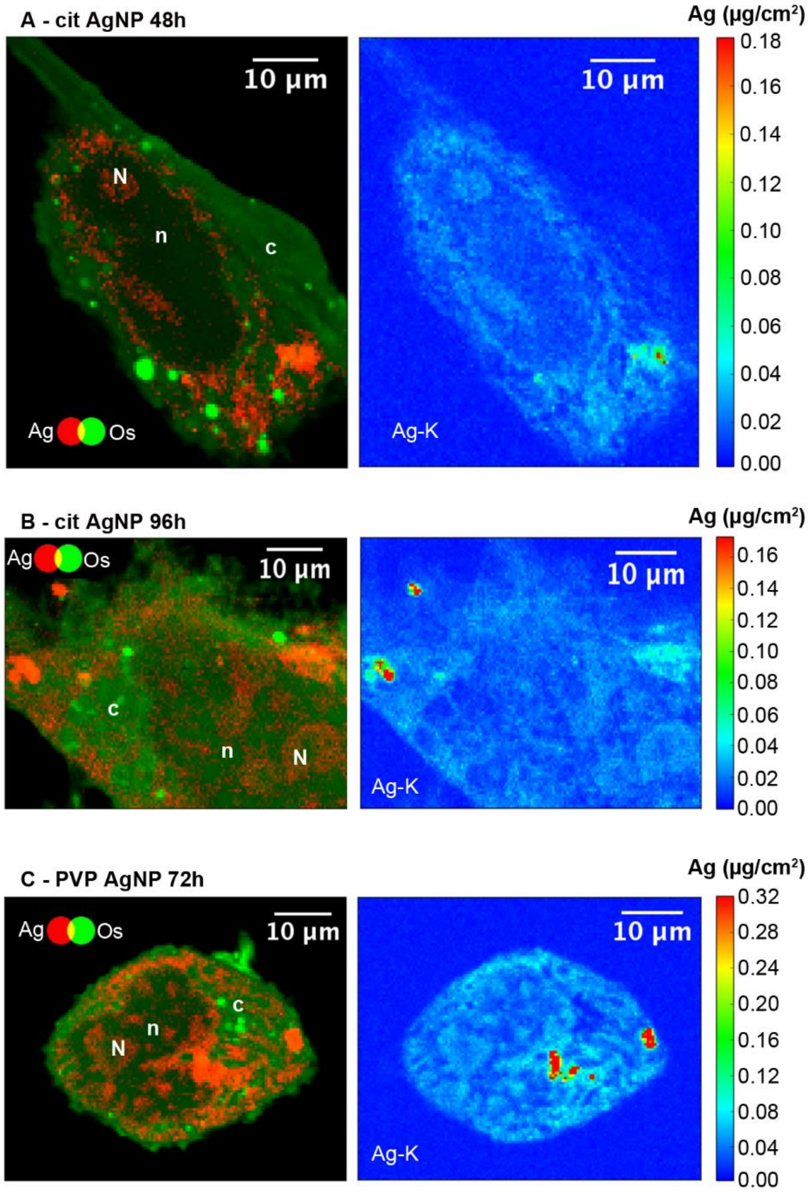


**Fig. S5.** Semi-thin sections of HepG2 cells analyzed by Nano-XRF. Two-color maps (Ag in red and Os in green) for 400 nm sections (left panels) and Ag areal density maps of the same regions (right panels) of cells exposed for, (A) 48h to 12.5 µM cit-AgNPs, (B) 96h to 12.5 µM cit-AgNPs, (C) 72 h to 25 µM PVP-AgNPs. Areal densities are expressed in µg/cm$^2$; pixel size is 100x100 nm$^2$. Scale bars correspond to 10 µm. c: cytoplasm; n: nuclei; N: nucleoli. The upper areal density limit in color bars is set to values that allow the visualization of the low areal density signal.

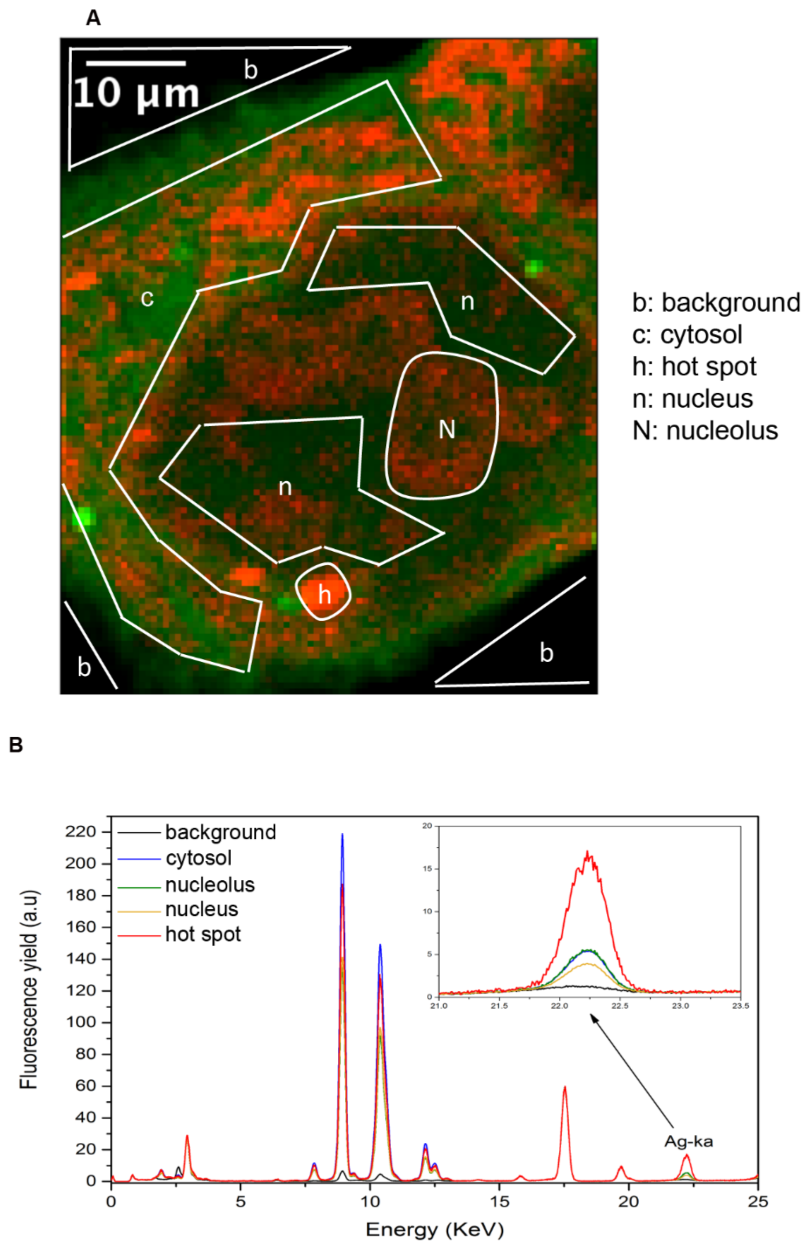


**Fig. S6.** Normalized XRF spectra from different sub-cellular areas. (A) two-color map (Ag in red and Os in green) for a 400 nm section of a HepG2 cell exposed for 72 h to 12.5 µM cit-AgNPs. Different areas in the map are defined as background (b), hot spots of Ag (h), cytoplasm (c), nucleus (n) and nucleolus (N). (B) normalized XRF spectra extracted from these different areas are presented with AgNP hot spots (red), cytosol (blue), nucleolus (green), nucleus (yellow) and background (black) regions. In the inset, a magnification of the Ag Kα emission region is shown.

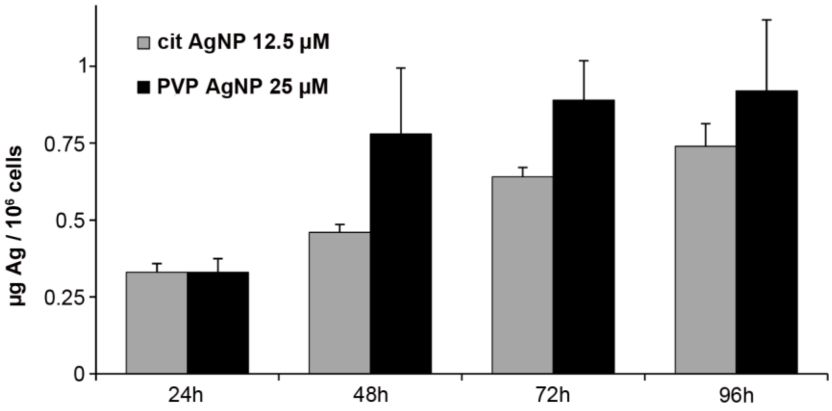


**Fig. S7.** Kinetics of cellular AgNP uptake. The amount of Ag per million cells was determined by ICP-AES after 24, 48, 72 and 96 hours of exposure to 12.5 µM cit-AgNPs or 25 µM PVP-AgNPs.

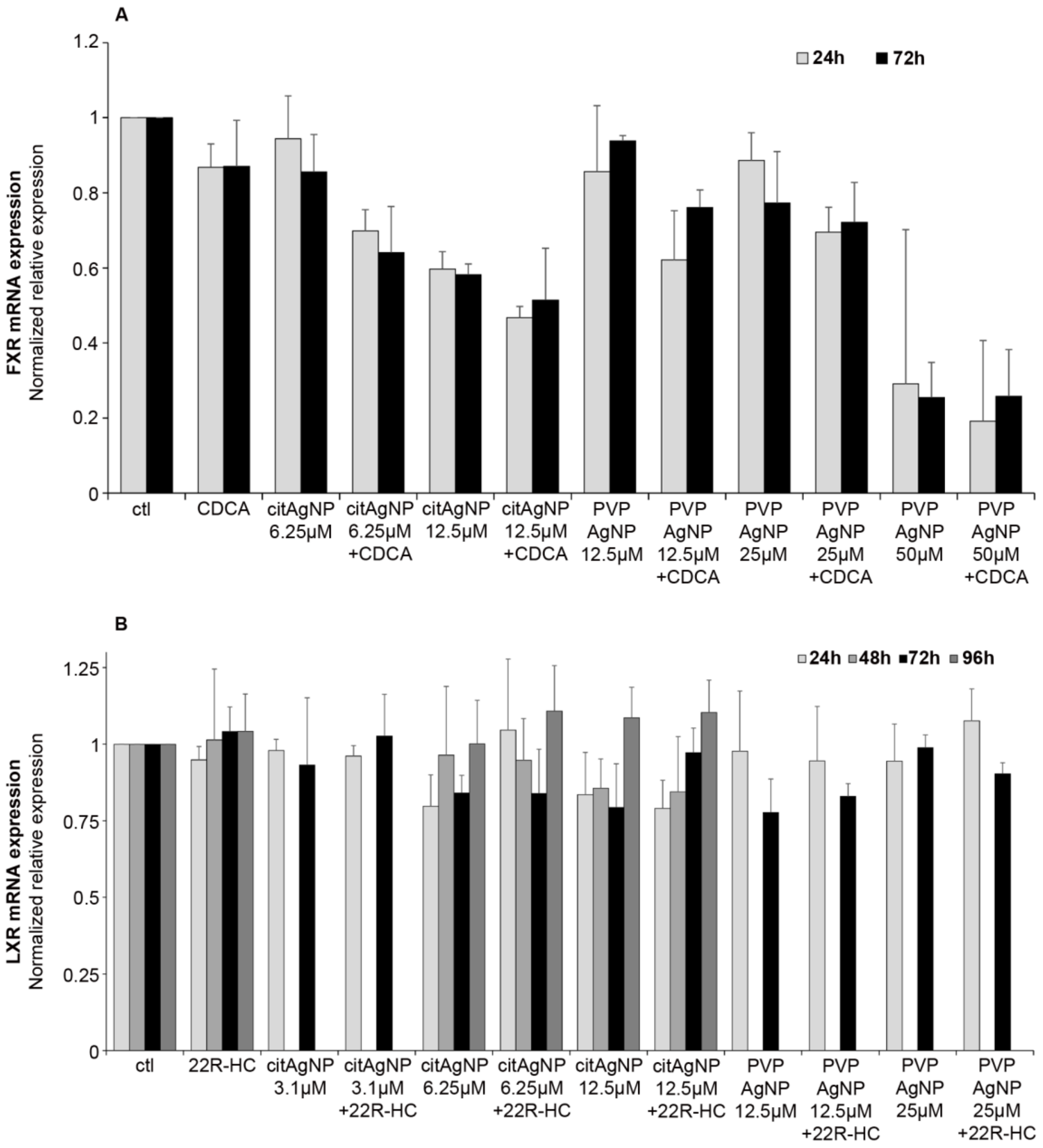


**Fig. S8.** Variations of nuclear receptor mRNA levels upon ligand addition and/or AgNP exposure. (A) FXR mRNA fold variation in HepG2 cells exposed to AgNPs and CDCA or DMSO. (B) LXRβ mRNA fold variation in HepG2 cells exposed to AgNPs and 22R-HC or ethanol. Results are expressed as the relative change in expression compared to the control (ctl), corresponding to cells unexposed to AgNPs. The results are expressed as means +/- standard error of the mean of at least three independent experiments.

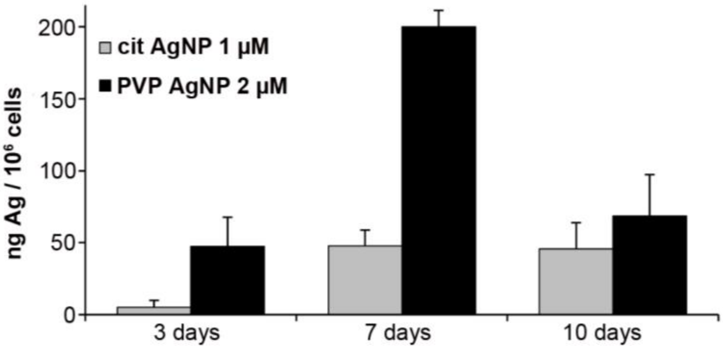


**Fig. S9.** Kinetics of cellular AgNP uptake under very low dose AgNP exposure. The amount of Ag per million cells was determined by ICP-AES after 3, 7 and 10 days of exposure to 1 µM cit-AgNPs or 2 µM PVP-AgNPs.

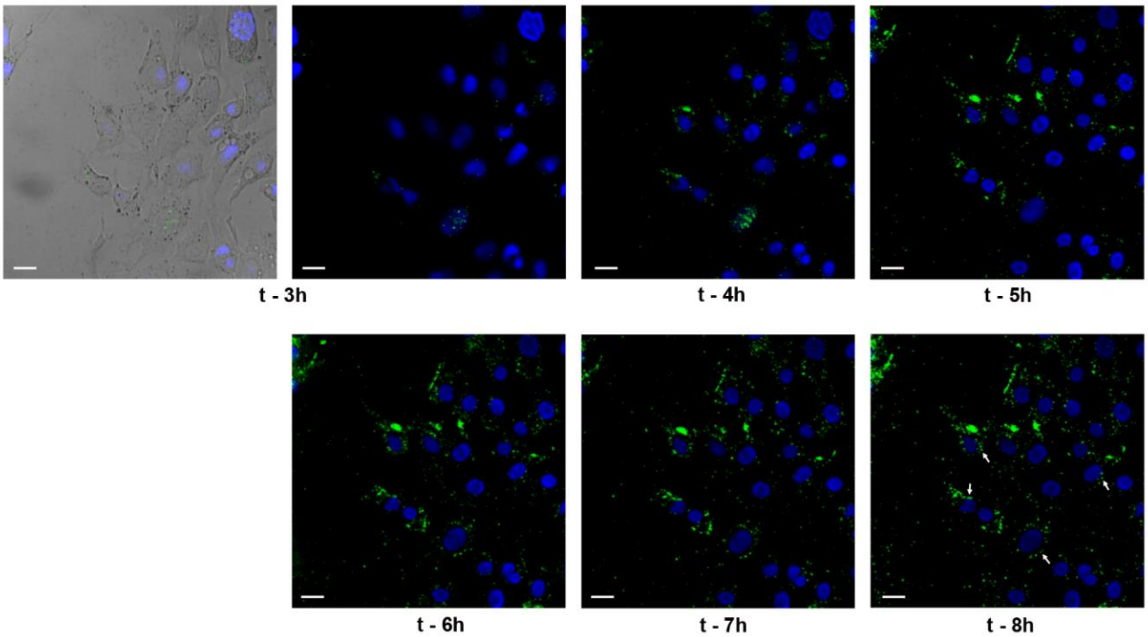


**Fig. S10.** Dynamics of cit-AgNPs in HepG2 cells. Confocal fluorescence and reflectance microscopy of HepG2 cells exposed to 12.5 µM of cit-AgNPs for the indicated times. Nuclei were labeled with Hoechst (blue) and AgNPs were observed in reflection mode (green). At t= 3 hours, a bright field image is also shown. The images correspond to the full field of the images shown in Fig. 4A. White arrows pinpoint particles close to nuclei. The scale bars represent 20 µm.

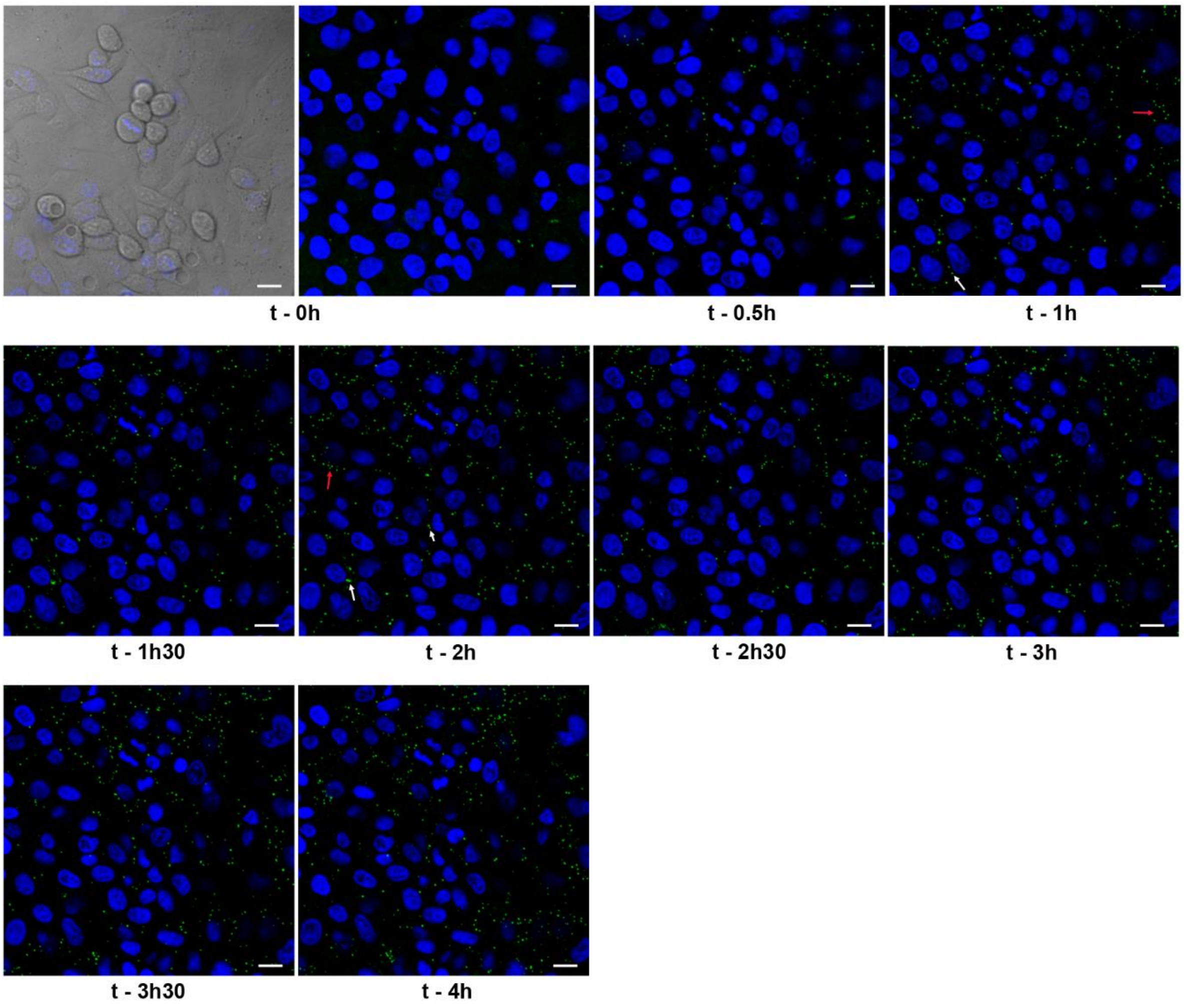


**Fig. S11.** Dynamics of PVP-AgNPs in HepG2 cells. Confocal fluorescence and reflectance microscopy of HepG2 cells exposed to 25 µM of PVP-AgNPs for the indicated times. Nuclei were labeled with Hoechst (blue) and AgNPs were observed in reflection mode (green). At t = 0 hour, a bright field image is also shown. The images correspond to the full field of the images shown in Fig. 4B and all the points in the kinetic. White arrows pinpoint particles close to nuclei and red arrows particles bound to the cell membrane. The scale bars represent 20 µm.

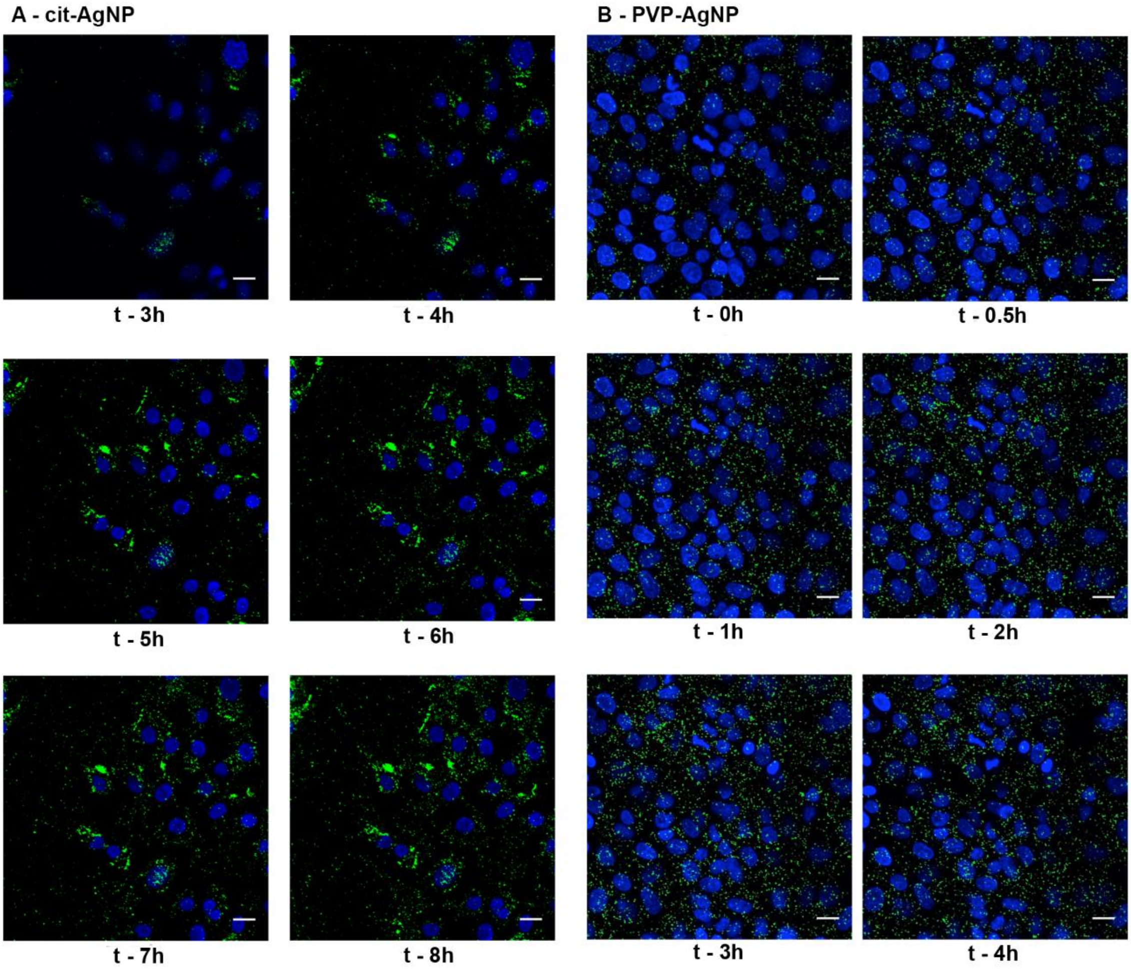


**Fig. S12.** Volume projections from confocal reflectance microscopy kinetics. Nuclei were labeled with Hoechst (blue) and AgNPs were observed in reflection mode (green). Images represent a maximum z-projection of all acquired planes, (A) cit-AgNPs, (B) PVP-AgNPs. The scale bar on all images is 20 µm.

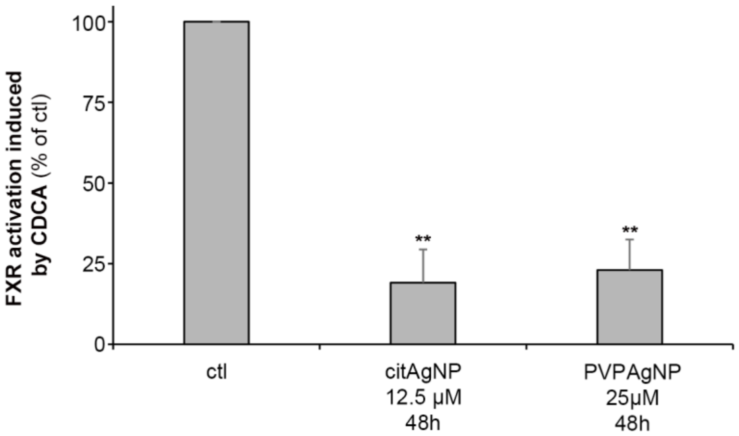


**Fig. S13.** Impact of AgNP exposure on FXR activity in hepatocyte spheroid. FXR activity followed by CDCA-induced BSEP expression. HepG2/C3a cells grown as spheroids were exposed or not to 12.5 µM cit-AgNPs or 25 µM PVP-AgNPs for 48 hours and to CDCA at 80 µM or DMSO during the last 17 hours. The RNA levels of BSEP were quantified in the different conditions using qPCR. 100% FXR activity was set using control conditions (unexposed to AgNPs) by calculating the ratio: amount of BSEP-RNA in ctl+CDCA / amount of BSEP-RNA in ctl without CDCA. This ratio (+CDCA/-CDCA) was measured for each condition and divided by the control ratio in order to determine the percentage of FXR activity in cells exposed to AgNPs (for details see experimental section). These experiments were performed three times independently. ** stands for data statistically different from the control with $p<0.01$.

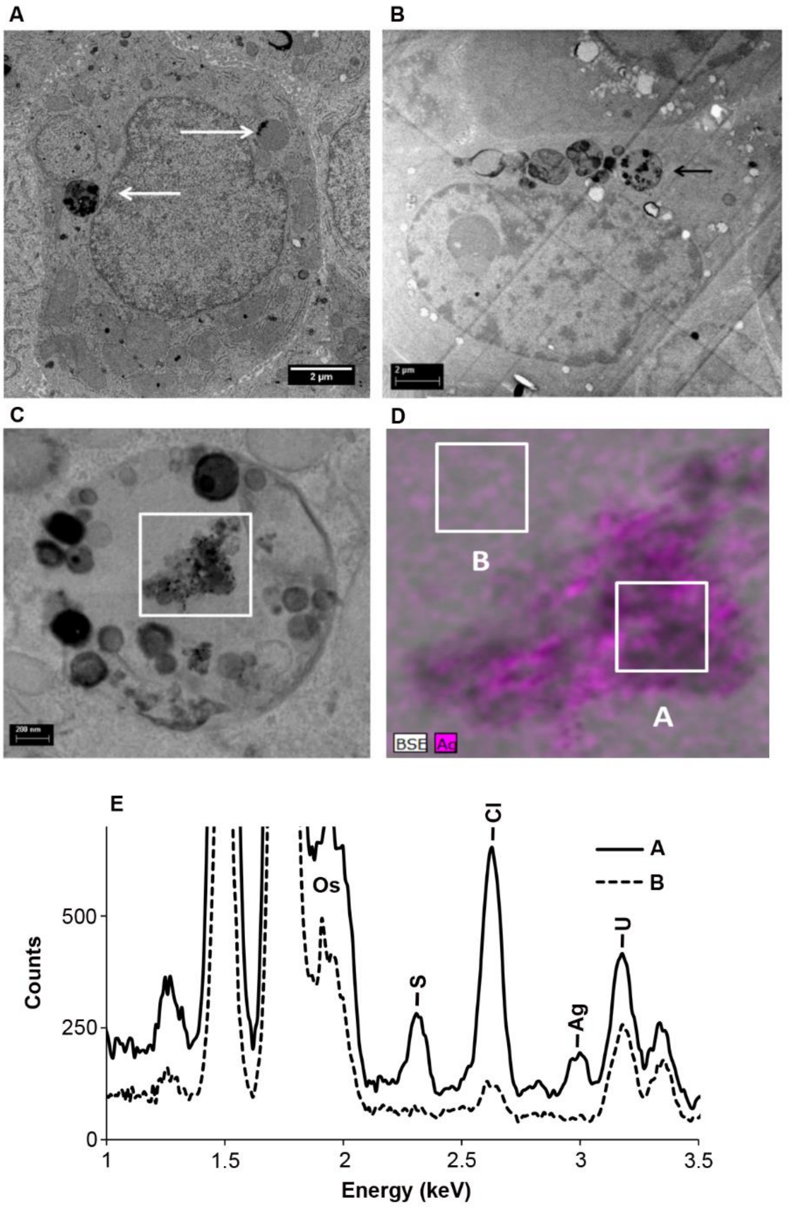


**Fig. S14.** Visualization and analysis of AgNP-containing vesicles close to nuclei. (A) and (B) STEM micrographs from 240 nm thin sections showing the presence of AgNP-containing vesicles close to nuclei (white and black arrows). Scale bars represent 2 µm. (C) Zoom in on the vesicle in (B) pointed by the black arrow. Scale bar represent 200 nm. (D) EDX map of the white box shown in (C) with Ag EDX map (pink) superimposed on the STEM map. (E) EDX spectra of two regions of the map shown in (D) noted A and B. The position of specific peaks of Os, S, Cl, Ag and U are highlighted in the spectra and proves the presence of Ag in the region A but not in the region B.

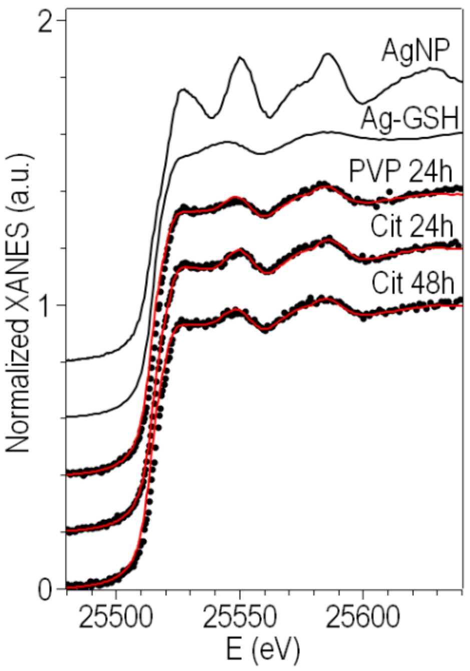


**Fig. S15.** Ag K-edge XANES of HepG2 cells exposed to AgNPs. Experimental spectra of reference compounds (solid black curves) and of cell pellets (black circles) exposed for 24 h to 12.5 µM cit-AgNPs or 25 µM PVP-AgNPs, or for 48h to 12.5 µM cit-AgNPs. The best-fitting curves obtained by linear combination of reference spectra are drawn in red over the corresponding experimental spectra.

### Supplementary tables

| | Hydrodynamic diameter (polydispersity index) | Zeta potential |
|---|---|---|
| PVP-AgNP | 89 nm (13.9%) | -6.8 mV |
| cit-AgNP | 160 nm (16.3 %) | -27.1 mV |

**Table S1. Characterization of AgNPs.** Diameter and polydispersity index (in parenthesis) of the main species observed in the dynamic light scattering distribution (in intensity) of cit-AgNPs and PVP-AgNPs in complete culture medium (Minimum Essential Medium + 10% Foetal Bovine Serum). The zeta potential was measured in storage buffer using a Malvern Nano ZS.

| Gene | Forward primer sequence | Reverse primer sequence |
|---|---|---|
| HPRT | ATGGACAGGACTGGACGTCTTGCT | TTGAGCACACAGAGGGCTACAATG |
| GAPDH | ATGGGGAAGGTGAAGGTCG | GGGGTCATTGATGGCAACAATA |
| MT1X | GCTTCTCCTTGCCTCGAA | TGACGTCCCTTTGCAGATG |
| ABCG5 | CTCCTACAGCGTCAGCCAC | AGGATGCACATGATCTGCCC |
| BSEP | ACATGCTTGCGAGGACCTTTA | GGAGGTTCGTGCACCAGGTA |
| FXR | GCATGCAGATCAGACCGTGA | TCAGTTTTCTCCCTGCATGACT |
| LXR-β | CAGAGCGCAAGCGAAAGAAG | GCTGAGCACGTTGTAGTGGA |
| 45S pre-rARN | ACCTGTCGTCGGAGAGGTT | GACGCGCGAGAGAACAGCA |
| HMOX1 | ATGACACCAAGGACCAGAGC | GTGTAAGGACCCATCGGAGA |

**Table S2.** Quantitative real time polymerase chain reaction primer sequences.

| Exposure conditions | AgNP fraction [%] | Ag-GSH fraction [%] | $R_{fit}$ |
|---|---|---|---|
| PVP-AgNP 24h | 24 ± 2 | 76 ± 2 | $1.8 \cdot 10^{-4}$ |
| cit-AgNP 24h | 30 ± 2 | 70 ± 2 | $2.3 \cdot 10^{-4}$ |
| cit-AgNP 48h | 23 ± 2 | 77 ± 2 | $2.3 \cdot 10^{-4}$ |

**Table S3.** Results of Linear Combination Fitting of Ag K-edge XANES spectra of HepG2 cells exposed to 12.5 µM cit-AgNPs or 25 µM PVP-AgNPs.

**Caption for Movie S1.** This movie shows, first, a complete stack of 2D micrographs and then segmentation of the different elements with nuclei in green, mitochondria in red, AgNPs in black (all AgNPs are inside vesicles as shown in Fig. 5B-C) and intercellular space in grey.
